# The p97-UBXD8 complex modulates ER-Mitochondria contact sites by modulating membrane lipid saturation and composition

**DOI:** 10.1101/2021.12.08.471763

**Authors:** Rakesh Ganji, Joao A. Paulo, Yuecheng Xi, Ian Kline, Jiang Zhu, Christoph S. Clemen, Conrad C. Weihl, John G. Purdy, Steve P. Gygi, Malavika Raman

## Abstract

The intimate association between the endoplasmic reticulum (ER) and mitochondrial membranes at ER-mitochondria contact sites serves as a platform for several critical cellular processes, in particular lipid synthesis. Enzymes involved in lipid biosynthesis are enriched at contacts and membrane lipid composition at contacts is distinct relative to surrounding membranes. How contacts are remodeled and the subsequent biological consequences of altered contacts such as perturbed lipid metabolism remains poorly understood. Here we show that the p97 AAA-ATPase and its ER-tethered ubiquitin-X domain adaptor 8 (UBXD8) regulate the prevalence of ER-mitochondria contacts. The p97-UBXD8 complex localizes to contacts and loss of this complex increases contacts in a manner that is dependent on p97 catalytic activity. Quantitative proteomics of purified contacts demonstrates alterations in proteins regulating lipid metabolism upon loss of UBXD8. Furthermore, lipidomics studies indicate significant changes in distinct lipid species in UBXD8 knockout cells. We show that loss of p97-UBXD8 results in perturbed contacts due to an increase in membrane lipid saturation via SREBP1 and the lipid desaturase SCD1. Aberrant contacts in p97-UBXD8 loss of function cells can be rescued by supplementation with unsaturated fatty acids or overexpression of SCD1. Perturbation of contacts and inherent lipid synthesis is emerging as a hallmark to a variety of human disorders such as neurodegeneration. Notably, we find that the SREBP1-SCD1 pathway is negatively impacted in the brains of mice with p97 mutations that cause neurodegeneration. Our results suggest that contacts are exquisitely sensitive to alterations to membrane lipid composition and saturation in a manner that is dependent on p97-UBXD8.

## Main

Contact sites between the ER and mitochondria allow for compartmentalization of biosynthetic reactions such as lipid synthesis, calcium transport and apoptosis among others^1–3^. ER- mitochondria contact sites (henceforth referred to as contacts unless specified otherwise) form when the membrane of these organelles come into close apposition (observed to be between 5 – 100 nm) without fusion^2, 4^. Contacts are stabilized by the interaction between tethering proteins that reside in opposing membranes; thus, the ability to regulate transient interactions between these tethers (through their abundance or post-translational modifications) enables the rapid formation and dissolution of contacts to meet cellular needs^5–9^.

Regulated protein degradation via the ubiquitin proteasome system is an efficient means to modulate contacts as numerous ubiquitin-reliant protein quality control mechanisms surrounding the ER and mitochondria can be co-opted to modulate the contact site proteome^10, 11^. The p97 AAA-ATPase is an abundant, evolutionarily conserved, ubiquitin-selective unfoldase that regulates multiple protein quality control pathways surrounding the ER and mitochondria. p97 is ideally positioned to regulate contacts by mediating the extraction and degradation of membrane-embedded tethers or contact resident proteins^12–14^. Importantly, specificity in p97-regulated pathways occurs via association with numerous adaptors that recruit p97 to specific substrates^15, 16^. We isolated contacts (also known as mitochondria-associated membranes, MAM) from HEK-293T cells and probed for p97 and select adaptor proteins. Percoll gradient centrifugation of crude mitochondria releases associated ER membranes allowing for the purification of ER-mitochondria contacts^17^. Using a known contact site-enriched protein as a marker (fatty acid coenzyme A ligase 4, FACL4) as well as markers for mitochondria (TOMM20) and ER (SEC61*β*), we found that the ER-localized p97 adaptors UBXD8 and UBXD2 (but not cytosolic UBXN1) are enriched at contacts and that p97 is also present in these fractions (Fig. 1a). To determine the role of p97 complexes at contacts we used a split luciferase reporter wherein the N-terminal fragment of luciferase is targeted to mitochondria and the C-terminal fragment to the ER using established targeting sequences^18^. Functional luciferase activity is reconstituted when the two organelles establish close range contacts and luciferase activity can be measured using the substrate, Enduren. We verified the functionality of the reporter system in wildtype cells by over-expressing receptor accessory protein 1 (REEP1)^18^, a contact tether and found a robust increase in contacts (Supplementary Fig.1a). To determine what role if any p97- adaptor complexes may have at contacts, cells were transfected with split luciferase cDNAs and siRNAs and luminescence was measured. Loss of p97 and UBXD8 resulted in an increase in contacts, whereas depletion of UBXD2 or another p97 adaptor UBXD7 had no impact (Fig. 1b, c and Supplementary Fig.1b). Hence even though UBXD2 localizes to contacts it does not regulate them and we focus on the role of UBXD8 for the remainder of the study. To verify the specificity of the phenotype, we expressed wildtype p97 or UBXD8 siRNA-resistant cDNAs and were able to rescue the phenotype (Fig. 1b, c). However, individual ATP catalytic site mutants in p97 were unable to rescue (Fig. 1c). To assess the role of individual domains in UBXD8, we expressed point mutants in the ubiquitin associated (UBA) or ubiquitin-X (UBX) domains that serve to bind ubiquitin and p97 respectively, as well as deletion of the UAS domain that has been reported to mediate the oligomerization of UBXD8^19^. Loss of these functional domains in UBXD8 prevented rescue of increased contacts (Fig. 1b, Supplementary Fig. 1c). Two additional assays were used to verify the role of p97-UBXD8 in regulating contacts. We measured contacts by measuring the extent of co-localization between the ER and mitochondria by confocal microscopy (Supplementary Fig. 1d, e), and we performed transmission EM (TEM) studies in wildtype and UBXD8 knockout (KO) cells generated by CRISPR-Cas9 gene editing (Fig. 1d-f). UBXD8 KO cells had a significant increase in ER tubules closely apposed to mitochondria (Fig. 1e, f and Supplementary Fig. 1e). No defects in overall mitochondria number or morphology were apparent in UBXD8 KO cells (Supplementary Fig. 1f).

**Fig. 1.**
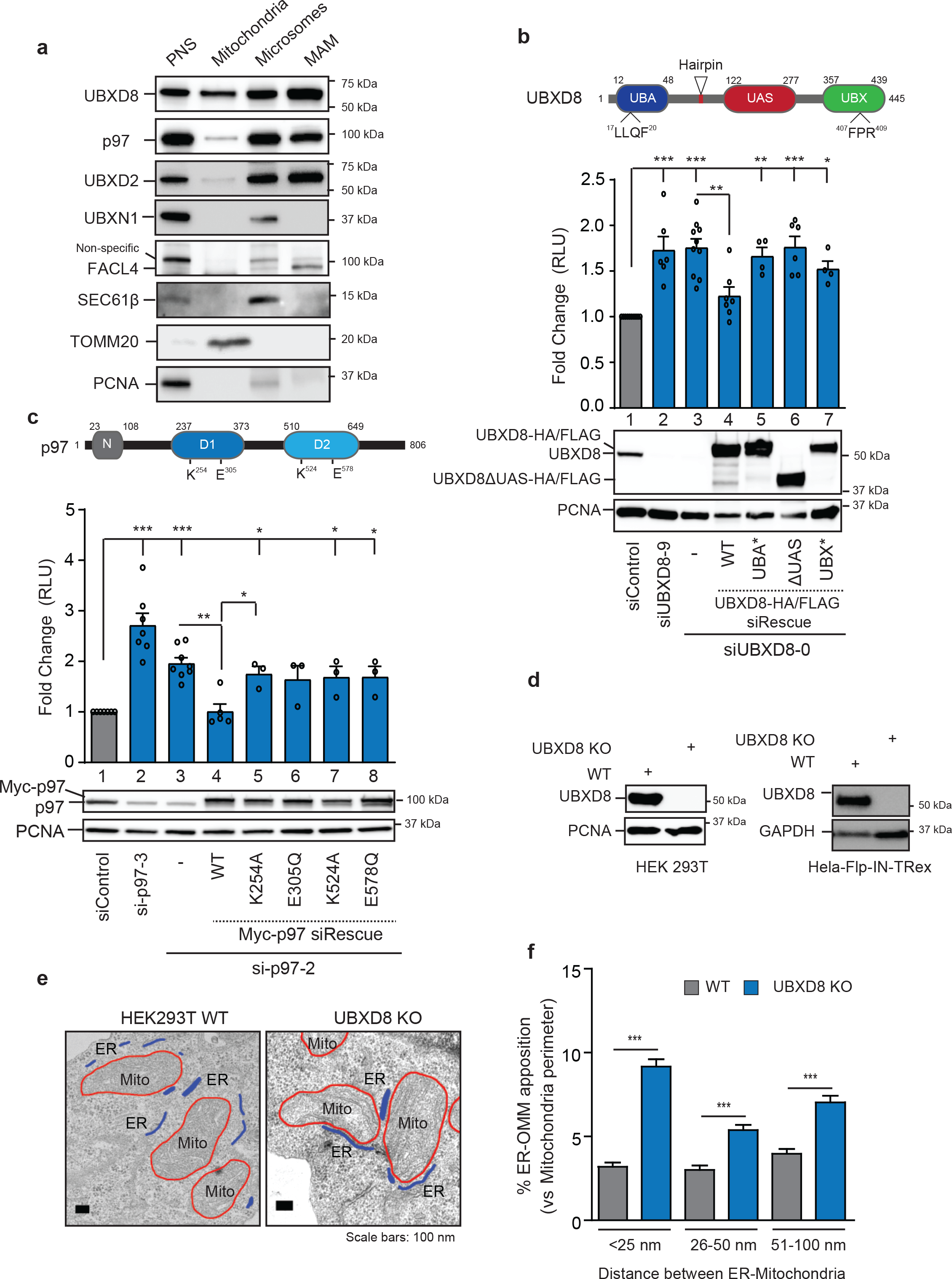
Loss of p97 and UBXD8 results in increased ER-mitochondria contacts. **a,** Immunoblot of indicated proteins from subcellular fractionation of HEK-293T cells, PNS: post- nuclear supernatant, MAM: Mitochondria associated membrane (*n* > 3 biologically independent samples). **b**, Top: Domain organization of UBXD8 and indicated mutations, Middle: Split luciferase assay to measure contacts in HEK-293T cells transfected with siRNAs to UBXD8 and indicated C-HA/FLAG siRNA-resistant rescue constructs, RLU: relative luminescence unit, Bottom: Immunoblot of indicated proteins. UBA: ubiquitin associated, UAS: upstream activating sequence, UBX: ubiquitin X (*n* > 3 biologically independent samples). **c**, Top: Domain organization of p97 and indicated mutations, Middle: Split luciferase assay to measure contacts in HEK-293T cells transfected with siRNAs to p97 and indicated N-Myc siRNA-resistant rescue constructs, RLU: relative luminescence unit, Bottom: Immunoblot of indicated proteins (*n* ≥ 3 biologically independent samples). **d**, Immunoblot of UBXD8 in CRISPR-Cas9 edited HEK-293T and HeLa-Flp-IN-T-Rex cells, KO: knockout. **e**, Representative transmission EM micrographs of wildtype and UBXD8 KO HEK-293T cells illustrating contacts between ER and mitochondria. **f**, Quantification of contact length between ER and mitochondria in each genotype from (**e**) (measurements are from *n* = 3 biological replicates with WT = 50 cells from 65 fields and UBXD8 KO = 53 cells from 71 fields). OMM: Outer mitochondrial membrane. Data are means ± SEM (*,**, \*\*\**P* < 0.05, 0.01, 0.0001 respectively, One-way ANOVA with Tukey’s multiple comparison test). Scale bar, 100 nm.

The p97-UBXD8 complex is critical for maintaining ER quality control by regulating ERAD and loss of this complex can cause ER stress due to deficits in ERAD^20^. To determine whether altered contacts observed upon depletion of p97-UBXD8 is due to increased ER stress, we treated cells with ER stressors tunicamycin and thapsigargin which cause protein misfolding in the ER. However, no impact on contacts was observed under these conditions (Supplementary Fig. 2a). Furthermore, we probed for the ER chaperone BiP which is induced upon ER stress and found no difference in BiP levels between wildtype and UBXD8 KO cells (Supplementary Fig.2b). Together with our findings suggest that the p97-UBXD8 complex has a novel role in regulating contacts independent of its role in ERAD.

To identify pathways altered in UBXD8 KO cells that may contribute to the contact site defect, we isolated contacts (MAMs) from triplicate wildtype and UBXD8 KO HEK-293T cells by biochemical fractionation and performed multiplexed, quantitative proteomics on the post-nuclear supernatant and MAM fractions using tandem mass tags (TMT) (Fig. 2a, b and Supplementary Fig. 3a-f, Supplementary Table 1)^21, 22^. We used two filtering criteria for downstream analysis: |log2 WT:KO ratio| > 1.0 and -log10 p value >1.5. The abundance of 23 proteins was enriched and 28 proteins was depleted in the MAM fraction of UBXD8 KO cells out of a total of 4499 quantified (Fig. 2b and Supplementary Fig. 3a). Putative contact site proteins identified in previous studies were present in our dataset validating our approach (Fig. 2b and Supplementary Fig. 3b, Supplementary Table 1) ^18, 23–25^. Furthermore, we identified significant enrichment of known p97- UBXD8 substrates such as squalene monooxygenase (SQLE), and HMG-CoA reductase (HMGCR) in UBXD8 KO cells (Fig. 2b and Supplementary Fig. 3g, h). Interestingly, proteins involved in lipid or cholesterol metabolism and lysosome function were also enriched in the UBXD8 KO contact proteome (log2 WT:KO ratio < -0.65 and -log10 p value >1.5) (Fig. 2b-d and Supplementary Fig.3c). In summary, our quantitative proteomic studies of the contact proteome in UBXD8 KO cells suggests that perturbation in the abundance of numerous enzymes linked to lipid biosynthesis may underlie the dysregulation in contacts.

**Fig. 2.**
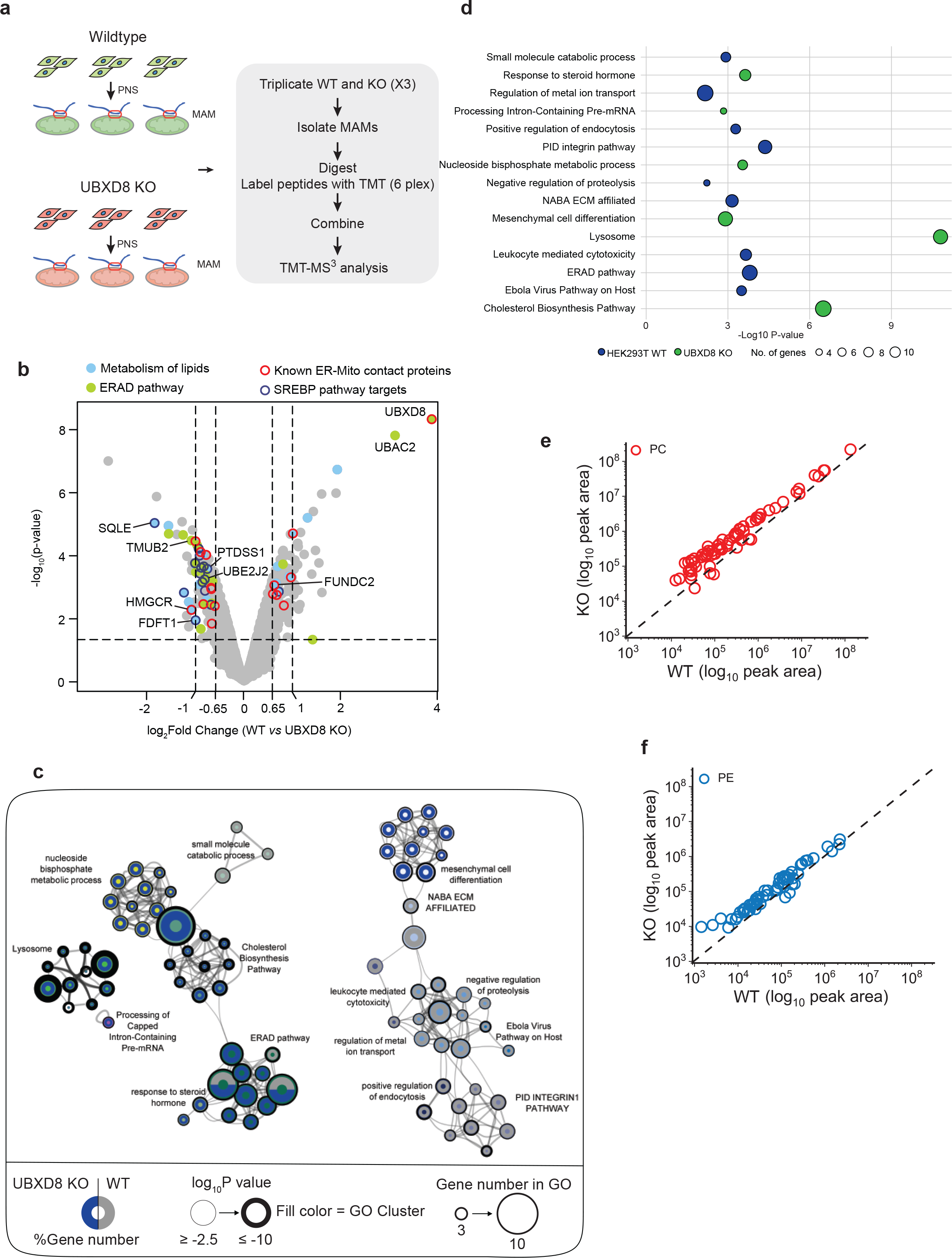
Quantitative proteomics and lipidomics identifies a role for UBXD8 in regulating lipid metabolism at contacts. **a,** Schematic of tandem mass tag (TMT) proteomic workflow from wildtype and UBXD8 KO HEK- 293T cells. PNS: post-nuclear supernatant, MAM: mitochondria associated membrane. **b**, Volcano plot of the (−log10-transformed *P* value *versus* the log2-transformed ratio of wildtype/ UBXD8 KO) proteins identified from MAM fractions of HEK-293T cells. *n* = 3 biologically independent samples for each genotype. *P* values were determined by empirical Bayesian statistical methods (adjusted for multiple comparisons) using the *LIMMA* R package; for parameters, individual *P* values and *q* values, see Supplementary Table 1. **c,** Network of differentially enriched terms shown as clustered functional ontology categories. Each node represents a functional ontology term enriched in the TMT data (**a, b**) as scored by Metascape^51^. Networks were generated using Cytoscape v3.8.2. Size of node represents number of genes identified in each term by gene ontology (GO). Grey and Blue donuts represent percent of genes identified in each GO term in wildtype or UBXD8 KO respectively. Node outline thickness represents −log10-transformed *P* value of each term. The inner circle color of each node indicates the corresponding functional GO cluster. **d**, Bubble plot representing significantly enriched GO clusters identified from TMT proteomics of MAM fractions in wildtype (blue) or UBXD8 KO (green) cells (**a-c**). Size of the circle indicates the number of genes identified in each cluster. **e-f**, Relative levels of Phosphatidylcholine (PC) (**e**) and Phosphatidylethanolamine (PE) (**f**) in HEK-293T WT and UBXD8 KO cells. PLs were measured by LC-MS/MS following normalization by total protein amount. Each dot in the plot represents the level of a PL in UBXD8 KO cells relative to its level in wildtype cells. The dashed line represents a relative level of 1 (e.g., the level in UBXD8 KO cells is equal to the level in wildtype cells). (*n* = 3 biologically independent experiments were performed, each with duplicate samples). Statistical analysis was performed on the log transformed relative fold change values (UBXD8 KO relative to WT) using independent *t* tests and Benjamini- Hochberg correction in R stats package (p-values are listed in Supplemental Table 1).

To better understand how cellular lipid metabolism may be impacted by loss of UBXD8 we measured the lipidome of wildtype and UBXD8 KO cells. We identified and quantitatively measured the relative levels of phospholipids (PLs) using liquid-chromatography high-resolution tandem mass spectrometry (LC-MS/MS). Our analyses examined the major classes of PLs found in membranes (and synthesized at contacts) including phosphatidylcholine (PC), phosphatidylethanolamine (PE), phosphatidylserine (PS), and phosphatidylinositol (PI). We also examined one-tailed lysophospholipids, which are metabolic byproducts of phospholipids (e.g., LPC, LPE, LPS, and LPI). We determined the concentration of 151 PLs in UBXD8 KO cells relative to their concentration in wildtype cells. In general, most PLs were more abundant in KO cells (Fig. 2e, f and Supplementary Fig. 4a-c, Supplementary Table 2). Approximately two-thirds of the two-tailed PC and PE and one-tailed LPC and LPE species were ≥2-fold more abundant in UBXD8 KO cells (Fig. 2e, f Supplementary Fig. 4a-d). The PC species that increased the most in the UBXD8 KO cells contained one or two double bonds among the fatty acyl tails (i.e., PC (44:1), PC (44:2), PC (46:1), PC (46:2), PC (48:2), PC (50:2), and PC (52:2)). MS/MS identification of the individual tails revealed that each of these lipids contained only saturated or monounsaturated fatty acids ranging in size from C16:0 to C32:1 and that lipids with one or fewer double bonds in each tail were increased the most by the loss of UBXD8 (Supplementary Fig. 4f and Supplementary Table 2). Phospholipids are synthesized via a diacylglycerol (DG) intermediate and DGs are metabolized to generate PLs and TGs. We therefore extended our lipidomics studies to measure DG and two additional neutral lipids: triacylglycerol (TG) and cholesteryl esters (CE) that are major components of lipid droplets whose biogenesis at the ER is regulated by UBXD8^26^. The relative concentration of most DGs were altered by less than 2-fold (Supplementary Fig. 4e). Of the ten DG species increased by ≥2-fold in UBXD8 KO cells, nine contained only saturated or monounsaturated tails ranging in size from C14:0 to C32:0. Similar to PC and PE, most TG species were ≥2-fold more abundant in KO cells relative to control cells (Supplementary Fig. 4e). The TG species whose abundance was enhanced the most by the loss of UBXD8 were also enriched in saturated and monounsaturated fatty acyl tails from C14:0 to C32:1. In contrast, most CE species was unaltered or slightly depleted by the loss of UBXD8 (Supplementary Fig. 4e). In summary, loss of UBXD8 shifts the lipidome to have a greater abundance of PC, PE, DG, and TG with saturated and monounsaturated fatty acyl tails demonstrating that UBXD8 is necessary for regulating lipid concentrations. The synthesis of these phospholipids occurs at contacts and their altered abundance in UBXD8 KO cells may contribute to defects in contacts^27^.

The significantly altered lipidome and related alterations in lipid biosynthetic enzymes prompted us to ask if these changes stem from the regulation of the sterol regulatory element-binding proteins 1 and 2 (SREBP1/2) pathway. p97-UBXD8 mediates the activation of ER-localized SREBPs by mediating ubiquitin-dependent membrane extraction and degradation of the SREBP negative regulator insulin induced gene 1 (INSIG1), in a sterol or fatty acid-dependent manner (Supplementary Fig. 5a)^28, 29^. A similar pathway for lipid sensing in *S.cerevisiae*, requires Cdc48p and Ubx2p (orthologs of p97 and UBXD8 respectively),^30^ and yeast lacking Ubx2p have more saturated cellular membranes due to loss of transcriptional activation of *Δ^9^* desaturase *ole1*^30^. We found that loss of p97 or UBXD8 resulted in a significant loss of SREBP1 activation and accumulation of the inactive ER-tethered form concomitant with stabilization of INSIG1 (Fig. 3a-d). Lipid desaturases are transcriptional targets of SREBP1 and were decreased in abundance at the transcript level in UBXD8 KO cells (Supplementary Fig.5b). This parallels the decreased protein abundance of the best characterized desaturases, SCD1 (*Δ^9^)* and FADS1 (*Δ^5^*) in UBXD8 KO or p97 depleted cells (Fig. 3a-d). In contrast, UBXD8 loss did not significantly impact SREBP2 activation or its downstream targets although this may be cell type or context specific (Fig.3a, b and Supplementary Fig.5c, d). Previous studies have reported that contacts have lipid raft-like properties and are enriched in cholesterol and sphingolipids^31^, thus the localization of cholesterol-sensing proteins to these sub-domains may be advantageous for rapid pathway activation. We find that SREBP1 and SCD1 are enriched at contacts isolated by biochemical fractionation (Fig. 3e). Notably, an increase in ER-tethered SREBP1 and a decrease in SCD1 were apparent in the MAM fractions of UBXD8 KO cells (Fig. 3f). Furthermore, over-expression of mature SREBP1a and c and to a lesser extent SREBP2, is sufficient to rescue contacts in p97-UBXD8 depleted cells (Supplementary Fig. 5e, f). Hence, defective SREBP1 activation underlies increased contacts in p97-UBXD8 depleted cells.

**Fig. 3.**
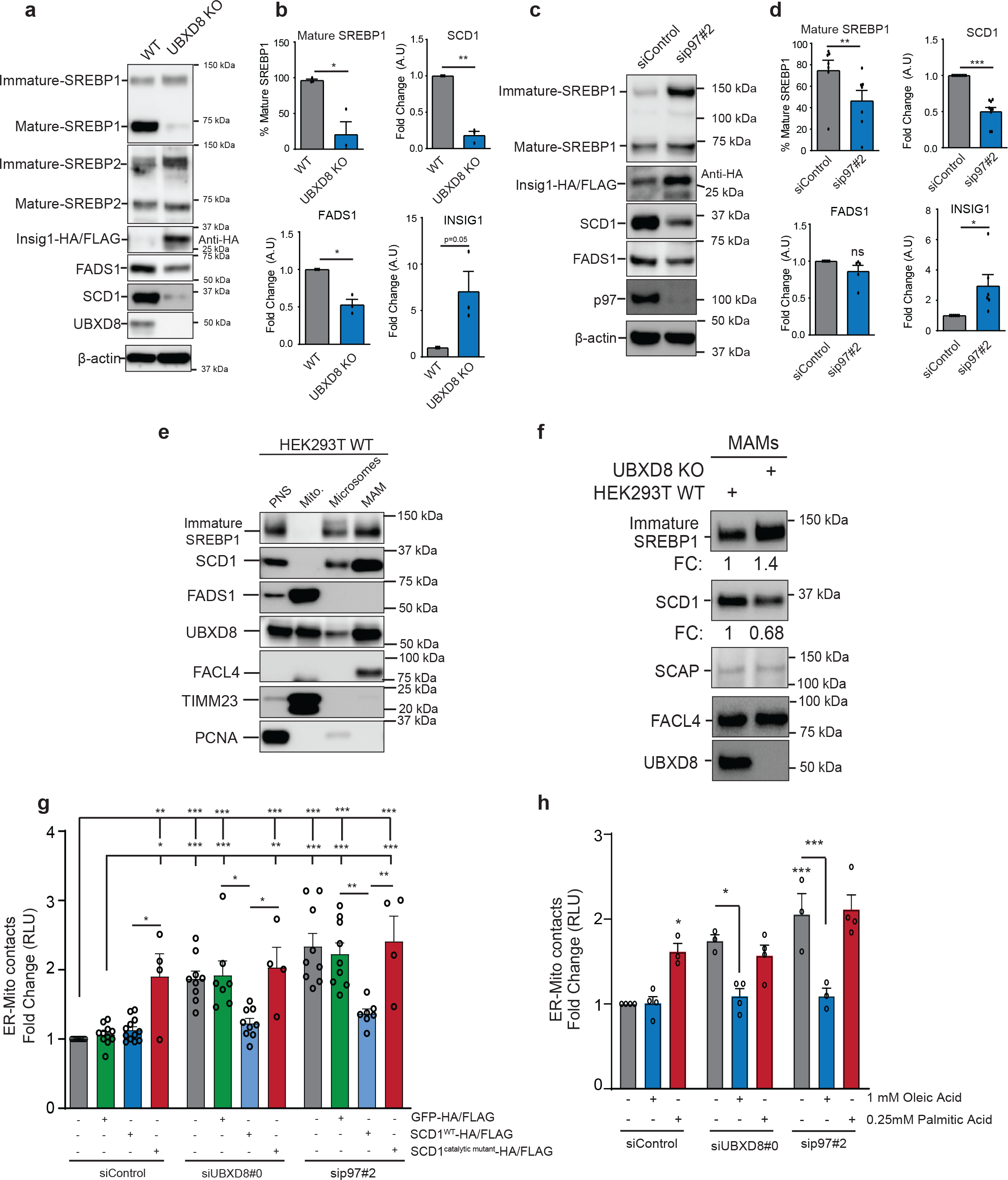
Loss of SREBP1 activation and SCD1 expression upon p97-UBXD8 depletion is responsible to contact defects. **a-d** Immunoblot and the corresponding band intensity quantifications of indicated proteins in the SREBP pathway in wildtype and UBXD8 KO HEK-293T cells (**a & b**), or p97-siRNA depleted cells (**c & d**). All samples were transfected with INSIG1-HA/FLAG due to lack of reliable antibodies to the endogenous protein. (*n* ≥ 3 biologically independent samples). **e**, Immunoblot of indicated SREBP pathway proteins from subcellular fractionation of HEK-293T cells, PNS: post-nuclear supernatant, MAM: mitochondria associated membrane. (*n* = 3 biologically independent samples). **f**, Immunoblot of indicated SREBP pathway proteins from subcellular fractionation of wildtype and UBXD8 KO HEK-293T cells, MAMs: mitochondria associated membrane. (*n* = 3 biologically independent samples). Corresponding fold changes (FC: UBXD8 KO *vs* WT) of SREBP1 and SCD1 normalized to FACL4 is shown. **g**, Split luciferase assay in HEK293T cells transfected with indicated siRNAs and wildtype or catalytically dead SCD1. GFP-HA/FLAG was transfected as a negative control. RLU: relative luminescence unit. (*n* ≥ 3 independent biological replicates). **h**, Split luciferase assay in HEK293T cells transfected with indicated siRNAs and treated with either monounsaturated oleic acid or saturated palmitic acid. RLU: relative luminescence unit. (*n* ≥ 3 independent biological replicates). Data are means ± SEM (*, **, \*\*\**P* < 0.05, 0.01, 0.0001 respectively. Paired *t* test (**b & d**), or One-way ANOVA with Tukey’s multiple comparison test (**g & h**).

The decrease in the abundance of lipid desaturases and our finding that UBXD8 KO cells have increased phospholipids with saturated or mono-unsaturated tails, prompted us to determine whether altering membrane lipid saturation perturbed contacts between the ER and mitochondria. FADS1 isoforms are localized to both ER and mitochondria^32^, therefore we focused on ER- localized SCD1^33^ as it was enriched at contacts (Fig. 3e). We treated wild type HEK-293T cells with the SCD1 inhibitor MF438^34^ and found that contacts between the ER and mitochondria increased in a manner that could be rescued by supplementing cells with monounsaturated oleic acid, the product of SCD1, (Supplementary Fig. 5g). We next evaluated whether re-expressing SCD1 in p97 or UBXD8-depleted cells rescued the increased contact site phenotype. Overexpression of wildtype SCD1 rescued ER-mitochondria contacts to wild type levels in p97- UBXD8 depleted cells (Fig. 3g, Supplementary Fig. 5h). In contrast, a catalytically inactive version of SCD1 was unable to rescue the phenotype (Fig. 3g, Supplementary Fig. 5h). Notably, overexpression of SCD1 catalytic mutant in wild type cells resulted in an increase in contacts suggesting that the resulting ordered lipid bilayers may impact contacts (Figure 3g). To further extend these findings, we asked if simply supplementing p97-UBXD8 depleted cells with unsaturated oleic acid (18:1), a precursor for the generation of polyunsaturated fatty acids in cells was sufficient to rescue contacts. Indeed, oleic acid but not saturated palmitic acid (16:0) rescued the contact defect in p97-UBXD8 depleted cells. Strikingly, palmitic acid alone increased contacts in wildtype cells (Fig. 3h).

Collectively, these results suggested that contacts are exquisitely sensitive to perturbations in lipid profiles within cellular membranes and that loss of p97-UBXD8 alters membrane lipid composition and saturation. We evaluated whether loss of p97-UBXD8 impacts lipid saturation globally within the cell. We used lipid-binding pyrene probes that insert into both disordered (unsaturated) and ordered (saturated) lipid bilayers throughout the cell and undergo monomer to excimer formation in a manner dependent on local membrane order. We measured the ratio of monomer to excimer fluorescence as an indicator of membrane order in cells treated with the p97 inhibitor CB-5083 or in UBXD8 KO cells. Loss of UBXD8 or inhibition of p97 resulted in more ordered cellular membranes (monomer: excimer ratio <1) compared to controls in two different cell lines (Fig.4a, Supplementary Fig.6a). Strikingly this phenotype can be reversed by incubating cells with oleic but not palmitic acid (Fig.4a, Supplementary Fig.6a). Thus, the local degradation of INSIG1 and subsequent activation of SREBP1 at contacts impacts lipid composition and saturation throughout the cell but significantly impact contacts due to their reliance on membranes for association. These findings prompted us to investigate whether the global increase in membrane order impacts other ER-organelle contacts. Analysis of TEM data from wildtype and UBXD8 KO cells demonstrated increased contacts between the ER and plasma membrane (Fig. 4b, c). Whether other ER-organelle contacts are similarly perturbed is under investigation. Notably, UBXD8 KO cells have an increase in membranous whorls containing concentric membrane layers, reminiscent of multilamellar bodies that function in lipid storage and secretion (Supplementary Fig. 6b). Interestingly, these membrane rich structures were also identified in yeast strains lacking Ubx2p ^30^ and may arise due to imbalances in lipid levels.

**Fig. 4.**
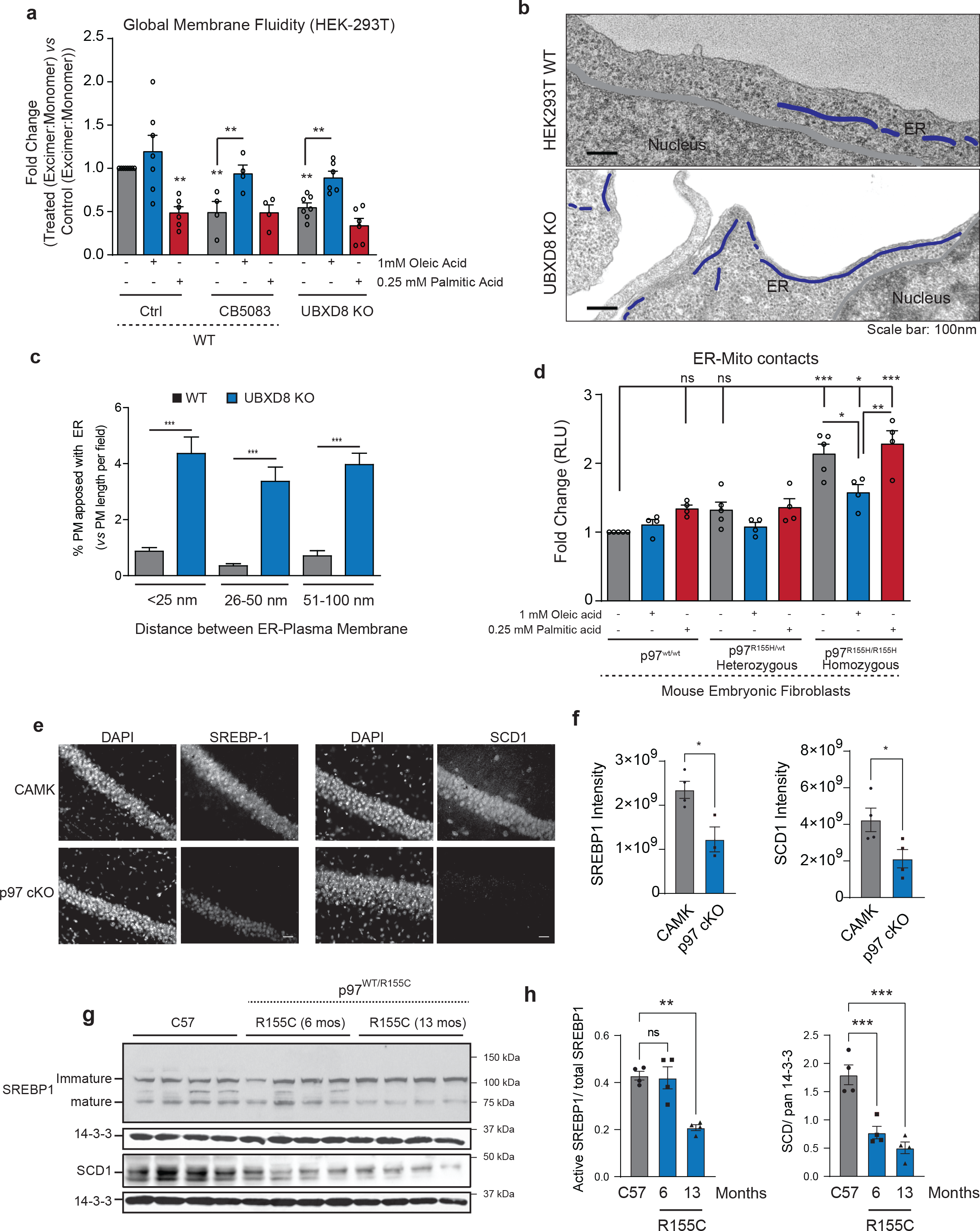
Supplementation of p97-UBXD8 depleted cells with oleic acid rescues contact defects. **a,** Global membrane fluidity was measured using a pyrene-based lipid probe in wildtype and UBXD8 KO HEK-293T cells. Wildtype cells were also treated with 5 *μ*M of the p97 inhibitor CB- 5083 for 4 hours. Cells were supplemented with indicated concentrations of oleic acid and palmitic acid for 4 hours. The fold change (Treatedexcimer:monomer *vs* Controlexcimer:monomer) of the ratio of excimer (Em. Max. 460nm) to monomer (Em max. 400nm) fluorescence is indicated. Fold changes < 1 indicate more ordered lipid bilayers relative to wildtype untreated control. (*n* ≥ 3 biologically independent samples). **b**, Representative transmission EM micrographs of wildtype and UBXD8 KO HEK-293T cells illustrating contacts between ER and plasma membrane. **c**, Quantification of contact length between ER and plasma membrane in each genotype from (**b**) (measurements are from *n* = 3 biological replicates with WT = 50 cells from 65 fields and UBXD8 KO = 53 cells from 71 fields). **d**, Split luciferase assay to measure contacts in mouse embryonic fibroblasts with heterozygous or homozygous p97 R155H mutation. Cells were supplemented with indicated concentrations of oleic acid and palmitic acid for 4 hours. (*n* ≥ 3 biologically independent samples). **e,** Representative SREBP1 and SCD1 staining from CA1 regions of 1 month-old control (CAMK2*α)* and p97 cKO mice (scale bar is 25 μm). **f,** Quantification of fluorescence intensity of images in (**e**). Individual points represent mean ROI intensity from each mouse, 3 or 4 animals per group. **g,** Representative immunoblot for SREBP1 and SCD1 from cortical brain lysates of 12-month-old control (C57), or 6- and 12-month-old p97^R155C/WT^ mice (n=4 for each group). Pan 14-3-3 was used as housekeeping control. **h,** Quantification of (**g**). The ratio of mature SREBP to total SREBP is shown. SCD1 intensities are normalized to 14-3-3 levels in each lane. Individual points represent each mouse, 4 animals per group. Data are means ± SEM (*, **, \*\*\**P* < 0.05, 0.01, 0.0001 respectively. Significance was analyzed by One-way ANOVA with Newman-Keuls multiple comparison test (**a**) or Tukey’s multiple comparison test (**c, d**) or Dunnett’s multiple comparison (**h**) or Student’s *t*-test (**f**).

Mutations in p97 cause several primarily neurodegenerative protein aggregation disorders. These include inclusion body myopathy with Paget’s disease of the bone and frontotemporal dementia (IBMPFD, also known as multi-system proteinopathy 1, MSP-1)^35^, amyotrophic lateral sclerosis (ALS)^36, 37^, Charcot Marie Type IIB^38^, among others. We investigated whether p97 disease-associated mutations altered contacts between the ER and mitochondria and perturbed the SREBP1-SCD1 pathway. We measured contacts in mouse embryonic fibroblasts heterozygous or homozygous for p97 R155H (a prevalent mutation observed in patients) and observed that cells with p97 R155H homozygous mutation had a significant increase in contacts that could be rescued with oleic acid but not palmitic acid (Fig. 4d).

Next, we evaluated the SREBP-SCD1 pathway in the brains of the two distinct p97 mouse models^39^. A conditional knockout of p97 (p97 cKO) in the cortex and hippocampus has recently been shown to develop cortical atrophy, neuronal loss and TDP43 inclusions reminiscent of frontotemporal dementia^39^. We stained for SREBP1 and SCD1 in the CA1 regions of one-month- old p97 cKO mice before neurodegeneration phenotypes are observed. Strikingly, p97 cKO mice had a significant decrease of SREBP1 and SCD1 immunoreactivity in the CA1 region compared to age-matched controls (Fig. 4e, f). To assess whether these defects were also present in a pathogenic context, we immunoblotted for SREBP1 and SCD1 in brain lysates from 6- and 13- month-old p97^R155C/WT^ mice^39, 40^. SREBP1 processing was significantly diminished at 13 months in p97^R155C/WT^ mice relative to controls (Fig. 4g, h). Similarly, loss of SCD1 protein levels was apparent at 6 months and continued to decline at 13 months in p97^R155C/WT^ mice (Fig. 4g, h). Collectively, our findings suggest that p97 mutations that cause disease may also have underlying lipid metabolism deficits that could contribute to disease pathology.

## Discussion

Here we have identified an unanticipated role for p97 and its ER-tethered adaptor UBXD8 in regulating ER-mitochondria contacts by perturbing membrane composition and fluidity in multiple cell types. We propose that altered lipid bilayers that arise upon loss of UBXD8 impact contacts in multiple ways: (1) preventing their dynamic association and disassociation due to loss of fluidity, and (2) negatively impacting the lateral movement of tethering proteins within membranes. UBXD8 is unique among p97 adaptors in its evolutionarily conserved and multifunctional roles in lipid sensing and metabolism^19, 26, 30, 41–43^. Our proteomic and lipidomic data from UBXD8 KO cells suggests that widespread changes in lipid metabolism is likely due to the inability to mobilize SREBP1 from the ER. Similar to the yeast *ubx2* deletion phenotype, we find decreased abundance of lipid desaturases, particularly SCD1 and complementation of p97-UBXD8 depleted cells with SCD1 rescues contact defects. However, given the significant changes in lipid profiles in UBXD8 KO cells, it is likely that SCD1 regulation is not the only mechanism at play. It remains to be determined whether UBXD8 also facilitates the degradation of contact tethers. We note that UBXD8 is present in pure mitochondrial fractions (Fig. 1a). Previous studies have observed dual localization of UBXD8 to ER and mitochondria^44, 45^ and recently, the yeast ortholog, Ubx2p was reported to localize to the outer mitochondrial membrane where it associates with the translocon in the outer mitochondrial membrane (TOMM) complex to clear stalled polypeptides in a p97-dependent manner^46^. Further studies are required to determine whether mammalian UBXD8 functions in an analogous manner on mitochondria. Aberrant contact sites are emerging as a common feature in the pathophysiology of a wide spectrum of human diseases ranging from diabetes to neurodegeneration^47–49^. We find that p97 mutations that cause proteinopathies also exhibit increased contacts and display significantly decreased levels of SREBP1 and SCD1. A recent report found that motor neurons from ALS patients with p97 mutations exhibited more contacts between the ER and mitochondria relative to controls^50^. Thus, altered organelles contacts and downstream lipid synthesis warrants further investigation in p97 associated diseases.

## Materials and Methods

### Cell culture, transfections, immunoprecipitations, and treatments

HEK293T, HeLa Kyoto Mouse embryonic fibroblasts (MEFs), and HeLa-Flp-IN-TREX (HFTs, gift from Brian Raught, University of Toronto) cells were cultured in Dulbecco’s modified Eagle’s medium, supplemented with 10% fetal bovine serum (FBS) and 100 units/mL penicillin and streptomycin. Cells were maintained in a humidified, 5 % CO2 atmosphere at 37°C. HeLa Kyoto wildtype cell line was a generous gift from Ron Kopito (Stanford University).

For siRNA transfections, HEK293T cells were trypsinized and reverse transfected with siRNAs. HeLa Kyoto, MEF, and HFT cells were trypsinized and seeded into a 12-well or 6-well dish 24 hours prior to siRNA transfections. In both reverse and forward transfections, the cells were transfected with 20 nM siRNAs using RNAiMax (Invitrogen) according to the manufacturer’s protocol. Cells were harvested 48-72 hours post transfection. For DNA transfections, HEK293T cells in 6 well plates were transfected with 0.75 *μ*g pcDNA3-Mit-NRluc91 and pcDNA3-CRluc92- ER, 0.75 *μ*g of UBXD8-C-HA/FLAG constructs, 1 *μ*g of N-Myc-p97 constructs, 1 *μ*g of p97-C-Myc, 1 *μ*g of SCD1-C-HA/FLAG constructs, 0.2-0.5 *μ*g of GFP-C-HA/FLAG, 0.75 *μ*g of pCIG construct, 0.75 *μ*g of REEP1, or 0.75 *μ*g each of 2X-FLAG-SREBP1a, 2X-FLAG-SREBP1c, 2X-FLAG- SREBP2 using Polyehtylenimine (PEI) at 1:4 DNA:PEI ratio and typically harvested 36-48 hours post transfection. HeLa Kyoto, MEF, and HFT cells were transfected with cDNA using Lipofectamine 2000 (Invitrogen) and the cells were harvested 36-48 hours post transfection. Cells were lysed in mammalian cell lysis buffer (50 mM Tris-Cl, pH 6.8, 150 mM NaCl, 0.5% Nonidet P-40, HALT Protease inhibitors (Pierce) and 1 mM DTT). Cells were incubated at 4°C for 10 min and then centrifuged at 14,000 rpm for 15 min at 4°C. The supernatant was collected, and protein concentration was estimated using the DC protein assay kit (Biorad). Protein G agarose (Pierce, Thermo Fisher scientific) and the indicated antibodies were used for immunoprecipitation at 4°C for 3-5 hours. Beads were washed 3-5 times in 1 ml mammalian cell lysis buffer and resuspended in 2X SDS sample buffer. Cells were treated with 1 *μ*M of Bortezomib, 5 *μ*M CB-5083, 5 *μ*g/ml of puromycin, 1 *μ*M SCD1 inhibitor (MF438), 1 mM Oleic acid, or 0.25 mM Palmitic acid for the indicated times (see figures for details). A full list of constructs used in this study can be found in Supplementary Table 3.

### Generation of CRISPR cell lines

The CRISPR-Cas9 gene editing system was used to generate UBXD8 knockout cell lines in HEK293T, and HFT cells. The guide sequence 5’ GTTAACCTGCAGGGGTCGTGA 3’ was cloned into the pX459 vector carrying the hSpCas9 and transiently transfected into HEK293T and HeLa-Flp-IN-TRex cells using Lipofectamine 3000 (Invitrogen) per the manufacturer’s protocol. 36 hours post-transfection the cells were selected with 1 *μ*g/ml puromycin for a further 24-36-hrs.

The surviving cells were then serially diluted into 96 well plates for clonal selection and expression levels were monitored by immunoblotting.

### Antibodies, siRNAs and Reagents

The p97 (10736-1-AP), UBXD8 (16251-1-AP), FACL4 (22401-1-AP), UBXD2 (21052-1-AP), HRD1 (13473-1-AP), Sec61*β* (51020-2-AP), Calnexin (10427-2-AP), UBXN1 (16135-1-AP), SREBP1 (14088-1-AP), SREBP2 (28212-1-AP), FADS1 (10627-1-AP), anti-GFP (66002-1-AP) and SCD1 (23393-1-AP) antibodies were from Proteintech Inc. The TIMM23 (H-8; sc514463), TOMM20 (F-10; sc17764), TOMM70 (A-8; sc390545), pan-ubiquitin (P4D1; sc8017), c-Myc (9E10; sc40), *β*-Actin (AC-15; sc69879), GAPDH (O411; sc47724), PCNA (PC10; sc56), UBE2J1 (B-6; sc377002), and UBE2S (C-1; sc390917) antibodies were obtained from Santa Cruz Biotechnologies. LC3B (D11; 3868S), and BiP (C50B12; 3177T) were from Cell Signaling Technologies. Derlin1 (A302-849A-T), p97 (A300-589A), and TEB4/MARCHIV (A304-171A-T) were from Bethyl laboratories. The following antibodies UBXD7 (PA5-61972; Invitrogen), anti-HA (16B12; MMS-101P, Covance), anti-FLAG (M2; F3165 Sigma Aldrich), INSIG1 (ab70784; Abcam) were used for immunoblotting. HRP conjugated anti-rabbit (W401B) and anti-mouse (W402B) secondary antibodies were from Promega. CB-5083 was a gift from Cleave Biosciences and Bortezomib was from Selleckchem. Palmitic acid (100905) is from MP Biomedicals and Oleic acid (270290050) is from Acros Organics. MF438 (569406) is from Millipore Sigma. All siRNAs were purchased from Ambion (Thermo Fisher Scientific). UBXD8-0 (s23260), UBXD8-9 (s23259), UBXD7-7 (s24997). UBXD2-1 (D-014184-03), UBXD2-2 (D-014184-04), HRD1-3 (D-007090-03), and HRD1-4 (D-007090-04) were purchased from GE Dharmacon. siControl (SIC001) was from Millipore Sigma. p97 siRNAs (2-HSS111263 and 3-HSS111264) were from Invitrogen (Thermo Fisher Scientific). p97 rescue constructs were previously published ^53^ and were resistant to siRNA # 2. UBXD8-C-HA/FLAG construct was previously published ^53^. The UBXD8 rescue constructs, including UBA* (^17^LLQF^20^ mutated to ^17^AAAA^20^), ΔUAS (deleted amino acids between 122-277), and UBX* (^407^FPR^409^ mutated to ^407^AAA^409^), were cloned using overlap PCR followed by Gibson assembly (NEB) cloning into pHAGE-C-HA/FLAG and were resistant to siRNA # 0. The SCD1-C- HA/FLAG WT and catalytic dead mutant (His^160^ His^161^ and His^301^ His^302^ mutated to Ala^160^ Ala^161^ and Ala^301^ Ala^302^) constructs were cloned using overlap PCR followed by Gateway cloning (Thermo Fisher Scientific) into pHAGE-C-HA/FLAG.

### Mitochondria-associated membranes (MAM) fractionation

MAMs were isolated as previously described ^54, 55^. Briefly, HEK293T or HeLa-Flp-IN-T-Rex cells were seeded into four 150 mm TC dishes. Cells were lysed in Homogenization buffer (225 mM mannitol, 75 mM sucrose, and 30 mM Tris-Cl, pH 7.4) using a Dounce homogenizer. The lysate was centrifuged three times at 600xg for 5 minutes to remove unlysed cells and nuclei resulting in post-nuclear supernatants (PNS). The cleared lysate was centrifuged at 7000xg to separate crude mitochondrial pellet and supernatant containing microsomes. The supernatant was cleared by centrifugation at 20,000xg for 30 minutes followed by microsome isolation using high-speed centrifugation at 100,000xg for 1 hour. The crude mitochondrial pellet was washed twice in homogenization buffer containing 0.1 mM EGTA at 7000xg and 10,000xg for 10 minutes. MAMs were isolated from crude mitochondria using 30% Percoll gradient centrifugation at 95,000xg for 1 hr in a swinging-bucket rotor. The banded MAM fraction was washed once with phosphate-buffered saline (PBS) before lysing in lysis buffer (50 mM Tris-Cl, pH 7.2, 150 mM NaCl, 2% SDS). The pure mitochondrial fractions were resuspended and washed in mitochondrial resuspension buffer (250 mM mannitol, 0.5 mM EGTA, 5 mM HEPES pH7.4). Mitochondrial membranes were solubilized using 0.5% (v/v) Digitonin. Protein concentrations for both soluble and pellet fractions were determined by DC protein assay kit (Biorad).

### Split luciferase assay to measure ER-Mitochondria contacts

Cells seeded a day prior in a 12-well plates were co-transfected with 0.75 *μ*g pcDNA3-Mit- NRluc91 and pcDNA3-CRluc92-ER (kind gift from Jeffrey A. Golden, Brigham and Women’s Hospital, Boston) using PEI at a 1:4 (DNA:PEI) ratio, or Lipofectamine 2000 (Invitrogen) as per manufacturer’s protocol. Media was changed after 6 hour and 18 hour later the cells are split into a clear bottom white 96-well plate with 50-100K cells per well. After 24 hours, 30 *μ*M of live-cell substrate Enduren (Promega) was added to cells and incubated for 2-3 hours in a 37°C incubator. The luminescence was measured using a SpectraMax iD3 multi-well plate reader. The luminescence measurements were normalized to the cell viability in each condition. Cell viability was measured using Cell Titer-Glo (Promega) according to the manufacturers’ instructions. Relative luminescence units (RLU) for each cell line were normalized to the DMSO treated samples to derive fold changes in RLU. Mean, standard error of means (SEM) and statistical significance were calculated by one way ANOVA with indicated post-hoc analysis using GraphPad Prism 5.01 (www.graphpad.com).

### Immunofluorescence and Microscopy

HFT cells stably expressing Sec61*β*-eGFP were grown on #1.5 cover slips in a 12 well plate and transfected with indicated siRNAs using RNAiMax. 48 hours post-transfection, cells were washed briefly in PBS and fixed with 4% paraformaldehyde at room temperature for 15 min. Cells were washed in PBS and permeabilized in ice-cold 100% methanol at -20°C for 10 min. Cells were washed three times in PBS and blocked in 2% BSA in PBS with 0.3% Triton X-100 for 1 hour. The coverslips were incubated overnight with the indicated antibodies in a humidified chamber. The cells were washed and incubated for a further hour with appropriate Alexa-Fluor conjugated secondary antibodies (Molecular Probes) for 1 hour in the dark. Cells were washed with PBS, nuclei were stained with Hoechst and mounted on slides. All images were collected using a Nikon A1R scan head with spectral detector and resonant scanners on a Ti-E motorized inverted microscope equipped with 60× Plan Apo NA 1.4 objective lens. The indicated fluorophores were excited with either a 405nm, 488nm or 594nm laser line. Images were analyzed using FIJI (<u>https:/t/imagej.net/Fiji</u>). Using a previously described method^52^, co-localized pixel analysis to quantify the ER-mitochondrial contact sites was performed using an ImageJ macro containing tubeness, colocalization highlighter, and isophotcounter plugins (Supplementary fig 2a).

### Lipid depletion and fatty acid supplementation

Cells were depleted of or supplemented with fatty acids as previously described^56–58^. Briefly, cells were treated with DMEM containing 0.5% lipid-depleted fetal calf serum (LDFCS; S5394, Sigma-Aldrich) for 24 hours. 500 mM Oleic acid in DMSO was used as a stock solution to prepare a working solution of 1 mM in DMEM containing 0.5% LDFCS. Cells were treated for 4 hr. 500 mM palmitic acid stock solution was prepared in 100% ethanol by heating to 70°C for 20 min. This stock solution, was used to prepare 0.25 mM palmitic acid solution in DMEM containing 0.5% LDFCS which was heated in a water-bath at 50°C for 2 hr. The 0.25 mM palmitic acid solution is cooled down to 37°C before adding to cells. Cell were incubated in palmitic acid solution for 4 hr. All working solutions were prepared immediately prior to treatment.

### Real time polymerase chain reaction (PCR)

Equal number of HEK293T WT or UBXD8 KO cells were seeded into a 6-well plate. The next day, total RNA was isolated as per manufacturer’s instructions using the PureLink RNA Mini kit (Thermo fisher). The purified RNA was quantified and an equal amount of RNA was used for cDNA preparation using iScript cDNA synthesis kit (Biorad). GAPDH was used as a housekeeping gene was used. Primer sequences used in this study can be found in Supplementary Table 3. Real time PCR was performed using the Powerup SyBr green master mix (Thermo Fisher). Data analyses were carried out using the 2^−ΔΔCt^ method.

### Membrane fluidity measurements

Cells were seeded in clear bottom black 96-well plate and treated with 5 *μ*M CB-5083, lipid depletion or lipid supplementation as described above. Membrane fluidity was measured using a membrane fluidity kit (Abcam, ab189819) as per manufacturer’s instructions. The assay uses a lipophilic, membrane embedding pyrenedecanoic acid probe which undergoes a spectral shift in emission from monomer (Em 400nm) to excimer (Em 460nm) based on local membrane fluidity upon excitation at 360nm. The excimer to monomer (Em 460nm / Em 400nm) ratio was calculated for each sample. Then fold changes of ratios (Treatedexcimer:monomer *vs* Controlexcimer:monomer) were deduced to provide a relative estimate of membrane fluidity compared to the wildtype untreated control. A fold change less than 1 indicates ordered membranes relative to control.

### Transmission Electron microscopy

Cells were fixed in 2.5% glutaraldehyde, 3% paraformaldehyde with 5% sucrose in 0.1 M sodium cacodylate buffer (pH 7.4), pelleted, and post fixed in 1% OsO4 in veronal-acetate buffer. The cells were stained *en bloc* overnight with 0.5% uranyl acetate in veronal-acetate buffer (pH 6.0), then dehydrated and embedded in Embed-812 resin. Sections were cut on a Leica EM UC7 ultra microtome with a Diatome diamond knife at a thickness setting of 50 nm, stained with 2% uranyl acetate, and lead citrate. The sections were examined using a FEI Tecnai spirit at 80KV and photographed with an AMT CCD camera. The images were analyzed manually for the ER-Mitochondrial and ER-PM contacts using FIJI (https:/t/imagej.net/Fiji). Briefly, the scale of image was set using Set Scale tool on ImageJ. Followed by measuring the length of ER, PM, or perimeter of mitochondria using freehand line tool. The percent of contact length was determined by taking the ratio of the length of ER, or PM (within contact distances of 25-100nm) to the perimeter of mitochondria. The data were analyzed using GraphPad Prism 5.01 for Windows, GraphPad Software, San Diego California USA, (www.graphpad.com).

### TMT-based proteomics

#### Sample preparation, digestion, and TMT labeling

The PNS and MAM fractions were isolated from HEK293T WT or UBXD8 KO cells. 100 *μ*g protein from each sample was precipitated using 15% (v/v) Trichloroacetic acid (TCA) followed by 100% Acetone washes. The protein pellets were resuspended in 200 mM N-(2-Hydroxyethyl)piperazine- N′-(3-propanesulfonic acid) (EPPS) (pH 8.5) buffer followed by reduction using 5 mM tris(2- carboxyethyl)phosphine (TCEP), alkylation with 14 mM iodoacetamide and quenched using 5 mM dithiothreitol treatments. The reduced and alkylated protein was precipitated using methanol and chloroform. The protein mixture was digested with LysC (Wako) overnight followed by Trypsin (Pierce) digestion for 6 hours at 37°C. The trypsin was inactivated with 30% (v/v) acetonitrile. The digested peptides were labelled with 0.2 mg per reaction of 6-plex TMT reagents (ThermoFisher scientific) (126, 127N, 127C, 128N, 128C, and 129N) at room temperature for 1 hour. The reaction was quenched using 0.5% (v/v) Hydroxylamine for 15 min. A 2.5 *μ*L aliquot from the labeling reaction was tested for labeling efficiency. TMT-labeled peptides from each sample were pooled together at a 1:1 ratio. The pooled peptide mix was dried under vacuum and resuspended in 5% formic acid for 15 min. The resuspended peptide sample was further purified using C18 solid-phase extraction (SPE) (Sep-Pak, Waters).

#### Off-line basic pH reverse-phase (BPRP) fractionation

We fractionated the pooled, labeled peptide sample using BPRP HPLC^59^. We used an Agilent 1200 pump equipped with a degasser and a detector (set at 220 and 280 nm wavelength). Peptides were subjected to a 50-min linear gradient from 5% to 35% acetonitrile in 10 mM ammonium bicarbonate pH 8 at a flow rate of 0.6 mL/min over an Agilent 300Extend C18 column (3.5 μm particles, 4.6 mm ID and 220 mm in length). The peptide mixture was fractionated into a total of 96 fractions, which were consolidated into 24 super-fractions^60^. Samples were subsequently acidified with 1% formic acid and vacuum centrifuged to near dryness. Each consolidated fraction was desalted via StageTip, dried again via vacuum centrifugation, and reconstituted in 5% acetonitrile, 5% formic acid for LC-MS/MS processing.

#### Liquid chromatography and tandem mass spectrometry

Mass spectrometric data were collected on an Orbitrap Lumos mass spectrometer coupled to a Proxeon NanoLC-1000 UHPLC. The 100 µm capillary column was packed with 35 cm of Accucore 150 resin (2.6 μm, 150Å; ThermoFisher Scientific). The scan sequence began with an MS1 spectrum (Orbitrap analysis, resolution 120,000, 350−1400 Th, automatic gain control (AGC) target 5 x10^5^, maximum injection time 50 ms). Data were acquired for 150 minutes per fraction. SPS-MS3 analysis was used to reduce ion interference^61, 62^. MS2 analysis consisted of collision-induced dissociation (CID), quadrupole ion trap analysis, automatic gain control (AGC) 1 x10^4^, NCE (normalized collision energy) 35, q-value 0.25, maximum injection time 60 ms), isolation window at 0.5 Th. Following acquisition of each MS2 spectrum, we collected an MS3 spectrum in which multiple MS2 fragment ions were captured in the MS3 precursor population using isolation waveforms with multiple frequency notches. MS3 precursors were fragmented by HCD and analyzed using the Orbitrap (NCE 65, AGC 3.0 x10^5^, isolation window 1.3 Th, maximum injection time 150 ms, resolution was 50,000).

### Data analysis

Spectra were converted to mzXML via MSconvert^63^. Database searching included all entries from the Human UniProt Database (downloaded: August 2018). The database was concatenated with one composed of all protein sequences for that database in the reversed order. Searches were performed using a 50-ppm precursor ion tolerance for total protein level profiling. The product ion tolerance was set to 0.9 Da. These wide mass tolerance windows were chosen to maximize sensitivity in conjunction with Comet searches and linear discriminant analysis^64, 65^. TMT tags on lysine residues and peptide N-termini (+229.163 Da for TMT) and carbamidomethylation of cysteine residues (+57.021 Da) were set as static modifications, while oxidation of methionine residues (+15.995 Da) was set as a variable modification. Peptide-spectrum matches (PSMs) were adjusted to a 1% false discovery rate (FDR)^66, 67^. PSM filtering was performed using a linear discriminant analysis, as described previously^65^ and then assembled further to a final protein-level FDR of 1% ^67^. Proteins were quantified by summing reporter ion counts across all matching PSMs, also as described previously^68^. Reporter ion intensities were adjusted to correct for the isotopic impurities of the different TMT reagents according to manufacturer specifications. The signal-to-noise (S/N) measurements of peptides assigned to each protein were summed and these values were normalized so that the sum of the signal for all proteins in each channel was equivalent to account for equal protein loading. Finally, each protein abundance measurement was scaled, such that the summed signal-to-noise for that protein across all channels equaled 100, thereby generating a relative abundance (RA) measurement.

Downstream data analyses for TMT datasets were carried out using the R statistical package (v4.0.3) and Bioconductor (v3.12; BiocManager 1.30.10). TMT channel intensities were quantile normalized and then the data were log-transformed. The log transformed data were analyzed with limma-based R package where p-values were FDR adjusted using an empirical Bayesian statistical. Differentially expressed proteins were determined using a log2 (fold change (WT *vs* UBXD8 KO)) threshold of > +/- 0.7.

### Gene ontology (GO) functional enrichment analyses of proteomics data

The differentially expressed proteins were further annotated and GO functional enrichment analysis was performed using Metascape online tool (http://metascape.org. The GO cluster network and protein-protein interaction network generated by metascape and the STRING database (https://string-db.org/), respectively, were imported into Cytoscape software (v3.8.2) to add required attributes (fold changes, p-values, gene number, and conditions) and prepared for the visualization. Other proteomic data visualizations were performed using the RStudio software (v1.4.1103), including hrbrthemes (v0.8.0), viridis (v0.6.1), dplyr (v.1.0.7), and ggplot2 (v 3.3.5).

### Lipidomics

#### Sample preparation, mass spectrometry, and identification

For each independent lipidomic experiment, HEK293T WT and UBXD8 KO cells were seeded in triplicate. Two of the three samples for each condition were used for lipidomics (i.e., lipids from duplicate samples were extracted and analyzed in parallel to determine technical variation). The third sample was used to determine the total protein concentration. Cells were washed with PBS, scraped into cold 50% methanol, centrifuged, and the cell pellets were frozen. Next, cells were resuspended in cold 50% methanol and transferred to glass vials. Chloroform was added and the mixture was gently vortexed and centrifuged at 1,000x g for 5 min at 4°C. Lipids were transferred to a clean glass vial using a glass Hamilton syringe. Lipids were extracted twice using chloroform prior to being dried under nitrogen gas. Samples were normalized according to protein concentration when resuspended in a 1:1:1 solution of methanol:chloroform:isopropanol prior to mass spectrometry (MS) analysis. The samples were stored at 4°C in an autosampler during data collection.

Lipids were identified and quantitatively measured using ultra high-performance liquid-chromatography high-resolution tandem MS/MS (UHPLC-MS/MS) as recently described^69, 70^. Separation of lipids was done by reverse-phase chromatography using a Kinetex 2.6 µm C18 column (Phenomenex 00F-4462-AN) at 60°C using a Vanquish UHPLC system (Thermo Scientific) and two solvents: solvent A (40:60 water-methanol plus 10mM ammonium formate and 0.1% formic acid) and solvent B (10:90 methanol-isopropanol plus 10mM ammonium formate and 0.1% formic acid). UHPLC was performed at a 0.25 ml per min flow rate for 30 min per sample, starting at 25% solvent B and ending at 100% solvent B as described. The column was washed and equilibrated between samples. Samples were run in a semi-random order where WT or UBXD8 KO samples were interspersed with blank samples. Lipids were ionized using a heated electrospray ionization (HESI) source and nitrogen gas and measured using a Q-Exactive Plus mass spectrometer operating at a MS1 resolution of either 70,000 or 140,000 and a MS2 resolution of 35,000. MS1 Spectra were collected over a mass range of 200 to 1,600 m/z with an automatic gain control (AGC) setting of 1e6 and transient times of 250 ms (70,000 resolution) or 520 ms (140,000 resolution). MS2 spectra were collected using a transient time of 120 ms and an AGC setting of 1e5. Each sample was analyzed using negative and positive ion modes. The mass analyzer was calibrated weekly. SPLASH LIPIDOMIX mass spectrometry standards (Avanti Polar Lipids) were used in determining extraction efficiencies and lipid quantitation. Quality control (QC) samples consisting of lipids extracted from the National Institute of Standards and Technology (NIST) Standard Reference Material 1950 Metabolites in Frozen Human Plasma which contains plasma pooled from 100 healthy donors were used in this study. In parallel to the samples, a control that lacked cells was used to determine any contaminants from the lipid extraction and measurement steps. Any lipids found in the no cell control were removed during analysis steps.

Lipids were identified and quantified using MAVEN^71^, EI-MAVEN (Elucidata), Xcalibur (ThermoFisher Scientific), and LipidSearch software (ThermoFisher Scientific). UHPLC retention time, MS^1^ peaks, and MS^2^ fragments were used to identify lipids. The lipid retention time, MS1 peak shape, isotopic distribution, and MS^2^ fragments were visually confirmed for all lipids reported in this study. Peak area was used to determined lipid abundance. Lipids were included if they were observed in 3-6 samples in both UBXD8 KO and WT cells. Missing values in a sample were not imputed. The fold change of each lipid in UBXD8 KO cells relative to its level in WT cells was used to test for statistical difference between UBXD8 KO and WT cells using independent t-tests and the Benjamini-Hochberg correction method to control for false statistical discovery. The following lipid classes were included in the analysis: cholesteryl esters (CE), diacylglycerol (DG), phosphatidylcholine (PC), phosphatidylethanolamine (PE), phosphatidylglycerol (PG), phosphatidylinositol (PI), phosphatidylserine (PS), and triacylglycerol (TG). Guidelines from the Lipidomic Standards Initiative were followed for lipid species identification and quantification, including consideration of isotopic patterns resulting from naturally occurring ^13^C atoms and isomeric overlap. The following MS^2^ information was used to confirm each lipid species: PC fragment of 184.073 (positive mode) and tail identification using formic adduct (negative mode); PE fragment of 196.038 or the tail plus 197.046 (negative mode) and neutral loss (NL) of 141.019 (positive mode); PG fragment of 152.996 plus the identification of the FA tails (negative mode) and NL 189.04 of [M+NH4]+ adduct (positive mode); PI fragment of 241.012 (negative) and NL 277.056 of [M+NH4]+ adduct (positive mode); PS NL of 87.032 (negative); DG and TG by NL of FA tails (positive mode); and CE fragment of 369.352 or neutral loss of 368.35 (positive).

### Mouse studies

C57BL/6 (stock No.: 000664) and p97^R155H/WT^ (B6;129S-Vcptm1Itl/J, Stock No: 021968) were purchased from Jackson Laboratory. p97^R155C/WT^ and p97 cKO (VCPFL/FL; CaMKIIa-Cre) were obtained as reported previously^39, 40^. All mice utilized in the study were on a C57BL/6 background. Both male and female mice were used in this study. Animal procedures were performed in accordance with protocols approved by the Animal Studies Committee at Washington University School of Medicine.

### Immunohistochemistry of tissue sections

Mice were anesthetized in an isoflurane chamber and perfused with PBS containing herapin. The whole brain was removed from the skull and fixed in 4% PFA overnight at 4 °C degrees, cut coronally into 40 micrometer sections, and stored in cryoprotectant solution at 4 °C degree for staining. Sections were first rinsed 3 times with TBS and then blocked with blocking solution for 30 minutes (5% normal goat serum with 0.1% Triton X-100 in TBS). Sections were stained with the primary antibody (SREBP1, 1:500, SCD1: 1:250 dilution) in TBS-0.1% Triton X-100 plus 2% normal goat serum at 4 °C overnight, followed by 3 washes with TBS. Sections were then incubated with the Alexa 488 and 555 tagged secondary antibodies (1:1000) for 2 hours at room temperature, followed by counterstaining with DAPI (1:1000) for 20 min. After three washes with TBS, the sections were mounted on the glass slides. True black (Biotium, NC1125051) was incubated with the sections for 5 minutes to quench the auto-fluorescence. Slides were cover- slipped using Prolong Gold mounting medium. Images were acquired using a Nikon Eclipse 80i fluorescence microscope. All images were taken with the same fluorescent settings and subsequently adjusted equally for brightness and contrast to ensure accurate pathology quantification. For CA1 regions, ROIs are drawn according to DAPI staining and the mean intensity was measured in each ROI by ImageJ. Background intensities (from three regions per image were subtracted.

### Immunoblot

Mouse cortex was lysed in RIPA buffer with protease inhibitor cocktails (PMSF and PIC) followed by sonication (two cycles of 30 seconds cycles at 50% power). The protein concentration is estimated by the BCA assay. Samples were loaded into 10% gel and transferred into nitrocellulose membrane. The membranes were blocked by 5% milk in PBS-0.2% Tween20 and incubated with the primary antibody in blocking solution overnight at 4 °C degrees. The membrane was then washed three times with PBS-0.2% Tween20 and incubated with a secondary goat anti-rabbit HRP antibody (1:5000) for 1 hour. Blot was rinsed three times with PBS-0.2% Tween20 and probed by a fresh mixture of ECL reagents at dark and then exposed by SYNGENE.

### Statistical analyses

For all experiments, N ≥ 3 biological replicates for each condition were examined. Fold changes, SEM, and statistical analyses were performed using GraphPad Prism version 5.01 for Windows, GraphPad Software, San Diego California USA (www.graphpad.com). The Statistical tests performed, SEM, and statistical significance values are mentioned in the figures and supplementary tables.

## Acknowledgements

We thank Peter Juo and Karl Munger for critical reading of the manuscript. We are grateful to Jeffrey Golden (Brigham and Women’s Hospital, Harvard Medical School) for split luciferase constructs, and the Whitehead Institute Electron Microscopy core for electron microscopy. We would also like to thank Jacob Klickstein for developing the ImageJ script for ER-mitochondria co- localization. This work is supported by the NIH grant GM127557 to M.R., funds from the University of Arizona Health Sciences and BIO5 Institute to J.G.P., and NIH grant AG031867 to C.C.W.

## Respective Contributions

R.G and M.R conceived the studies. R.G performed all experiments except for lipidomics studies (that were performed by Y.X. and J.G.P. and analyzed by Y.X., I.K. and J.G.P.) and mouse immunohistochemistry and immunoblotting (that were performed by J.Z). J.P and S.P.G assisted with proteomic studies. M.R wrote the manuscript with input from R.G and J.G.P and C.C.W.

## Competing Interests

The authors declare no conflicts of interest.

## Request for reagents

Please contact the corresponding author, M.R for reagent requests.

## Data availability

All raw proteomic and lipidomic data will be made available on public servers upon acceptance of the manuscript. Any other data is available from the corresponding author upon request.

## Code availability

The ImageJ macro to quantify ER-mitochondria contact sites will be made available on Github upon manuscript acceptance.

## Supplementary Figure Legends

**Supplementary Fig. 1.**
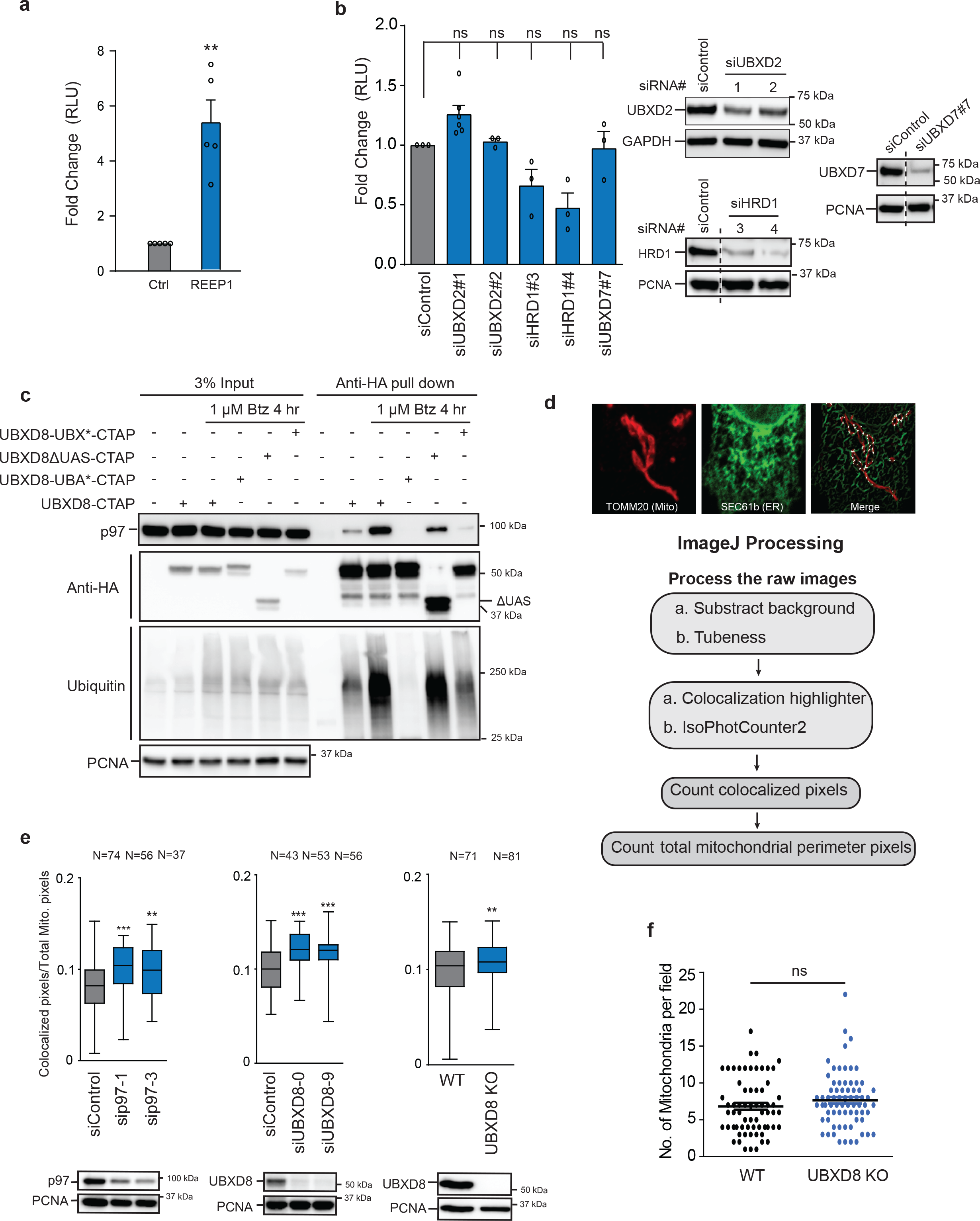
Loss of p97 and UBXD8 results in increased ER-mitochondria contacts. **a,** Split luciferase assay to measure contacts in HEK-293T cells transfected with REEP1 (*n* > 3 biologically independent samples). **b**, Left: Split luciferase assay to measure contacts in HEK- 293T cells transfected with indicated siRNAs. (*n* ≥ 3 biologically independent samples). Right: Immunoblot of HEK 293T cells transfected with indicated siRNAs. **c,** Affinity purification of indicated UBXD8-HA/FLAG constructs transiently expressed in HEK-293T cells. Immunoblots of whole cell lysates and affinity purifications probed with anti-HA, anti-p97, anti-PCNA, and ubiquitin antibodies (*n* = 3 biologically independent samples). **d**, Top: Representative confocal image of HeLa-Flp-IN-TRex cells stably expressing Sec61*β*-eGFP (green, ER) and stained for endogenous TOMM20, (red, mitochondria). Bottom: ImageJ image analysis pipeline for the quantification of contacts between ER and mitochondria^52^. **e**, Quantification of ER- mitochondria contacts in cells transfected with indicated siRNAs (left and middle panel) or in UBXD8 KO HeLa-Flp-IN-TRex cells (right panel) using assay in (**d**). Bottom panels show immunoblots for knockdown efficiency. (*n* = 3 biologically independent samples) N: numbers of cells analyzed in each condition. (Quartiles represent the upper 75th percentile and the lower 25th percentile. The line inside the box represents the median. Whiskers indicate distribution of data from minimum to maximum in a condition.). **f**, Quantification of number of mitochondria per field from transmission electron microscopy of wildtype and UBXD8 KO cells. (Measurements are from *n* = 3 biological replicates with WT = 50 cells in 65 fields and UBXD8 KO = 53 cells in 71 fields). Data are means ± SEM (*, **, \*\*\**P* < 0.05, 0.01, 0.0001 respectively. One-way ANOVA with Tukey’s multiple comparison test (**b & e** (left and middle panel)), Paired *t*-test (**a**) or Unpaired *t* test with Welch’s correction (**e** (right panel) & **f**).

**Supplementary Fig. 2.**
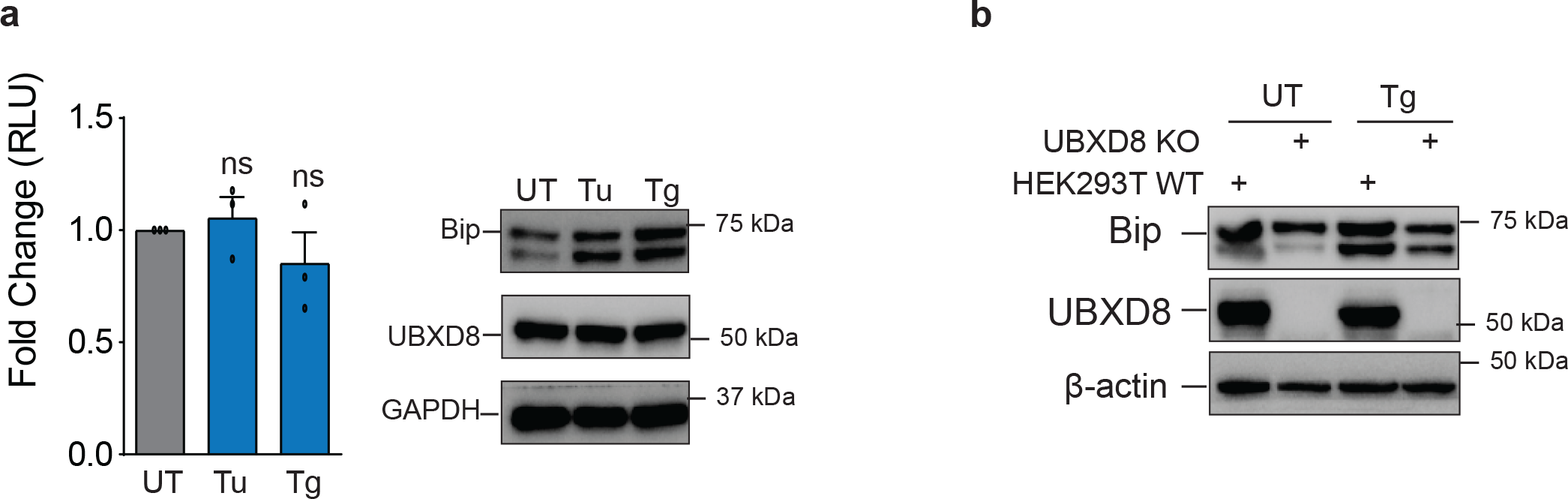
ER stress does not alter ER-mitochondria contacts. **a,** Split luciferase assay to measure contacts in HEK-293T cells treated with 2.5 µM Tunicamycin (Tu), 1.5 µM Thapsigargin (Tg), for 2 hours. (*n* = 3 biologically independent samples). Immunoblot of HEK-293T cells treated with Tu and Tg and probed for the indicated proteins. **b**, Immunoblot of HEK-293T wildtype and UBXD8 KO cells treated with 1.5 *μ*M Thapsigargin (Tg) for 2 hours, UT: Untreated. (*n* = 3 biologically independent samples). Data are means ± SEM (ns: not significant). One-way ANOVA with Tukey’s multiple comparison test (**a**).

**Supplementary Fig. 3.**
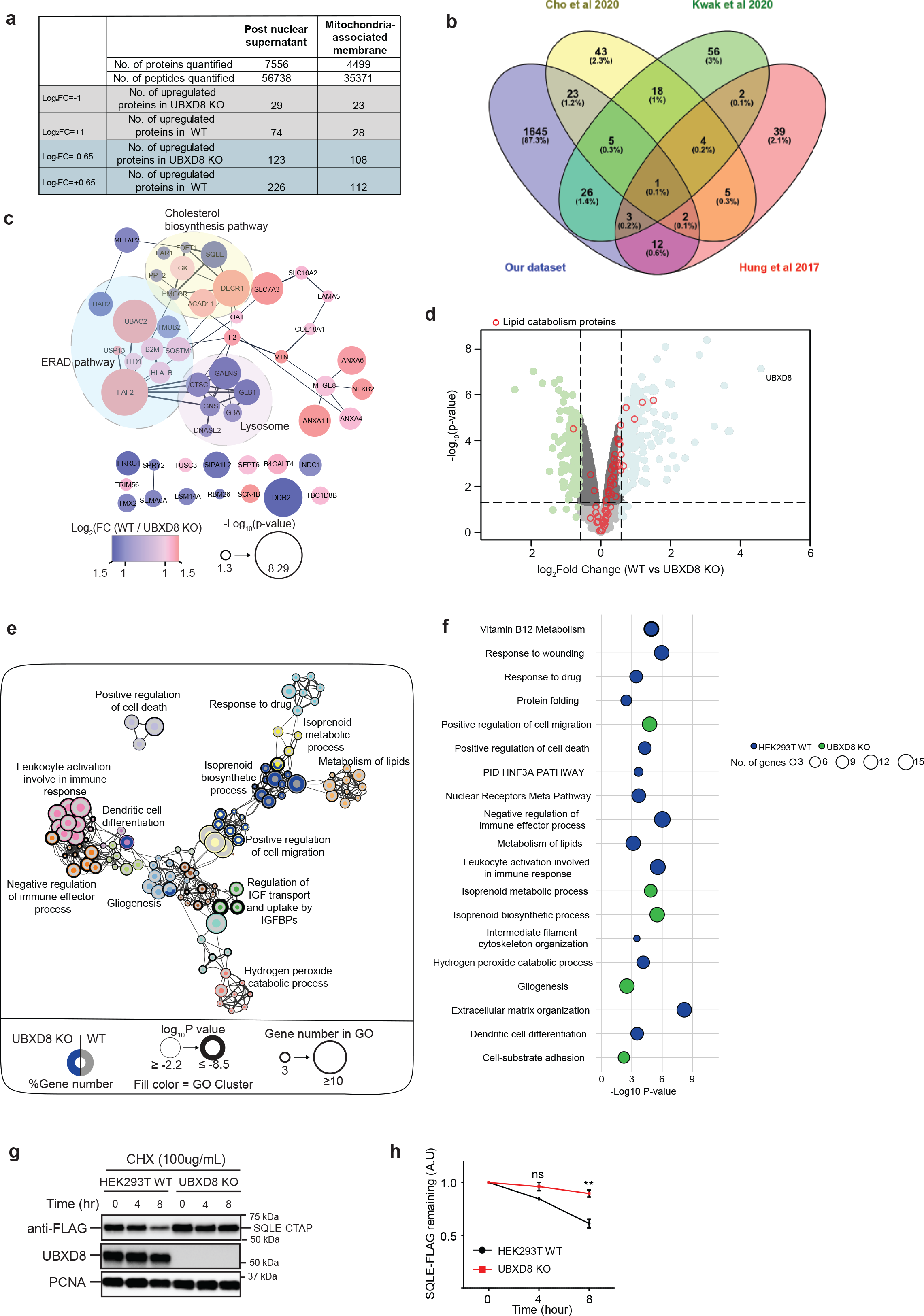
Quantitative proteomics of wildtype and UBXD8 KO contact proteome. **a,** Table depicting number of proteins and peptides quantified in post-nuclear supernatant and mitochondria associated membrane fractions identified by proteomics in wildtype and UBXD8 KO cells. Number of proteins up- or downregulated at log2- fold change (FC) (wildtype/ UBXD8 KO) ± 0.65 and ±1 is indicated. **b**, Venn diagram depicting overlap of our dataset with other putative mitochondria associated membrane proteins identified by proteomics^18, 23–25^ **c**, Protein-protein interaction network of differentially expressed proteins from MAM fractions of HEK-293T cells involved in ERAD, cholesterol biosynthesis and lysosome function shown as clustered functional categories. Protein associations were determined using STRING database with score ≥ 0.4. Each node represents a protein belonging to enriched GO clusters as scored by Metascape. Size of node represents −log10-transformed *P* value and color of node represents log2- fold change (FC) (WT / UBXD8 KO). **d**, Volcano plot of the −log10-transformed *P* value *versus* the log2-transformed ratio of wildtype/ UBXD8 KO proteins identified in the post-nuclear supernatant of HEK-293T cells. *n* = 3 (each genotype) biologically independent samples. *P* values were computed by empirical Bayesian statistical methods (adjusted for multiple comparisons) available in *Limma* R package; for parameters, individual *P* values and *q* values, see Supplementary Table 1. **e**, Network of differentially enriched functional ontology terms shown as clustered functional ontology categories. Each node represents a functional ontology term enriched in the TMT data (**d**) as scored by Metascape and networks generated using Cytoscape v3.8.2. Size of node represents number of genes identified in each term by gene ontology (GO). Grey and Blue donuts represent percent of genes identified in each GO term in wildtype or UBXD8 KO respectively. Node outline thickness represents −log10-transformed *P* value of each term. The inner circle color of each node indicates the corresponding functional GO cluster. **f,** Bubble plot representing significantly enriched GO clusters identified from TMT proteomics of post-nuclear fractions in wildtype (blue) or UBXD8 KO (green) cells (**d-e**). Size of the circle indicates the number of genes identified in each cluster. **g**,**h**, Squalene epoxidase (SQLE) half-life measurements in wildtype and UBXD8 KO HEK 293T cells. FLAG-SQLE was transiently expressed, and cells were treated with 100 *μ*g/mL cycloheximide for the indicated times. Samples were resolved on SDS-PAGE for immunoblots (**g**) and levels of SQLE were quantified and normalized to loading control PCNA (**h**). (*n* = 3 biologically independent samples). Data are means ± SEM (*, **, *** *P* < 0.05, 0.01, 0.0001 respectively. One-way ANOVA with Tukey’s multiple comparison test (**h**).

**Supplementary Fig. 4.**
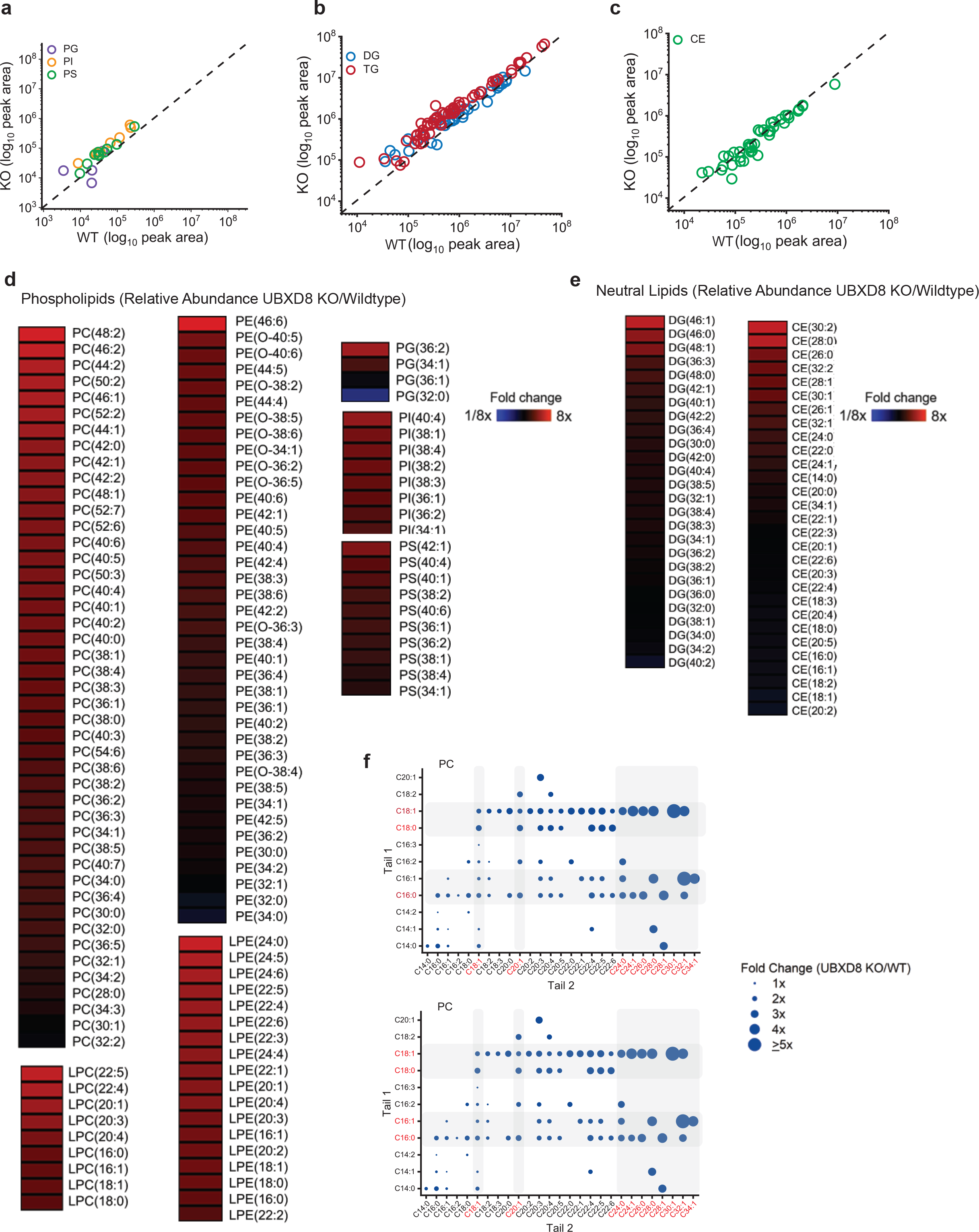
Loss of UBXD8 alters cellular lipidome with increased abundance of saturated fatty acid tail containing phospholipids. **a**, Relative levels of phospholipids (PLs), namely PG, phosphatidylglycerol; PI, phosphatidylinositol; and PS, phosphatidylserine in HEK-293T WT and UBXD8 KO cells. PLs were measured by LC-MS/MS following normalization by total protein amount. Each dot in the plot represents the level of a PL in UBXD8 KO cells relative to its level in wildtype cells. The dashed line represents a relative level of 1 (e.g., the level in UBXD8 KO cells is equal to the level in wildtype cells). (n = 3 biologically independent experiments were performed, each with duplicate samples). **b**, Relative levels of DG, Diacylglycerol; TG, Triacylglycerol in HEK-293T WT and UBXD8 KO cells. The DG and TG is quantified and visualized similarly as in **a**. (n = 3 biologically independent experiments were performed, each with duplicate samples). **c**, Relative levels of CE, Cholesteryl esters in HEK-293T WT and UBXD8 KO cells. The CE is quantified and visualized similarly as in **a**. (*n* = 3 biologically independent samples). **d**-**e**, Changes in the relative levels of PLs (**d**) including lysophopholipids (LPC, Lysophosphatidyl choline; LPE, Lysophosphatidyl ethanolamine; LPS, Lysophosphatidyl Serine; and LPI, Lysophophatidyl Inositol) and (**e**) Neutral lipids (DGs, TGs, and CEs) in HEK-293T WT and UBXD8 KO cells were quantified. The averaged relative fold changes were log2 transformed and visualized as a heatmap. (*n* = 3 biologically independent experiments were performed, each with duplicate samples). **f**, Dot plots representing fold change of PC and PE lipids in UBDX8 KO cells relative to WT cells and fatty acid tails determined by MS/MS. The two tails of the lipid are shown and organized so that tail 1 (y-axis) is the shorter tail and tail 2 (x-axis) is the longer tail. The labels tail 1 and tail 2 do not represent their stereospecific number (sn). Grey boxes indicate the increase in saturated and mono-unsaturated tails in UBXD8 KO cells. Statistical analysis was performed on the log2 transformed relative fold change values (UBXD8 KO relative to WT) using independent t tests and Benjamini-Hochberg correction in R stats package (p-values are listed in Supplemental Table S3).

**Supplementary Fig. 5.**
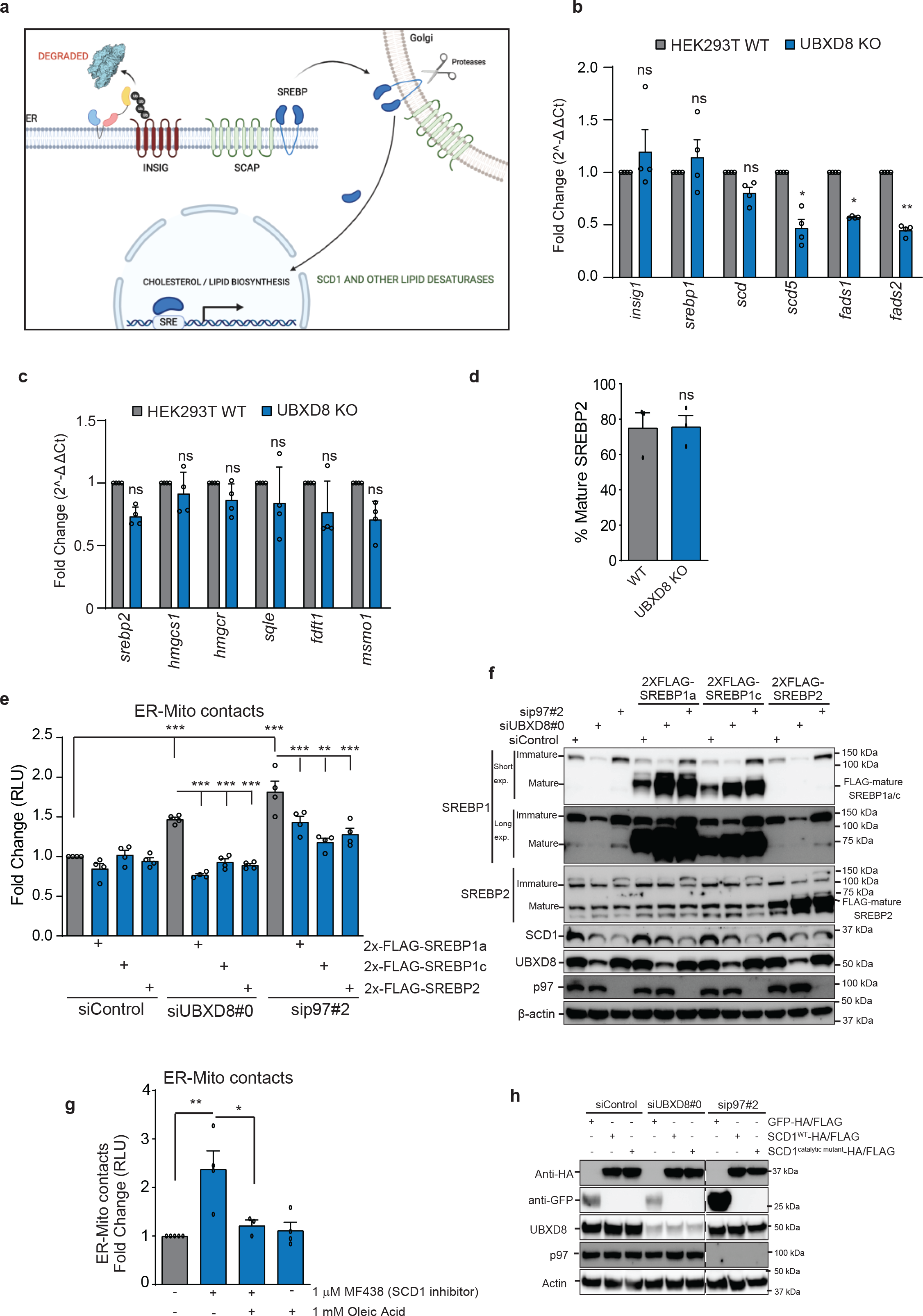
Diminished SREBP pathway activation in UBXD8 KO cells. **a,** Schematic of SREBP pathway activation. In cholesterol-replete conditions, SCAP-INSIGs- SREBPs are in an inactive tripartite complex in the ER membrane. Cholesterol depletion triggers a conformational change in SCAP releasing it from INSIGs and enabling the transport of SCAP- SREBPs to the Golgi. Here SREBPs are cleaved sequentially by site 1 and site 2 proteases to release the active transcription factor. INSIGs are ubiquitylated and extracted for degradation from the membrane by p97-UBXD8. **b-c,** Real-time quantitative PCR of SREBP1 target genes including lipid desaturases (**b**), and SREBP2 target genes (**c**). **d,** Band intensity quantifications of mature SREBP2 in wildtype and UBXD8 KO HEK-293T cells corresponding to **Fig 3a** (*n* ≥ 3 biologically independent samples) **e,** Split luciferase assay to measure contacts in HEK-293T cells transfected with siRNAs to UBXD8 or p97 and indicated 2X-FLAG-tagged mature SREBP1a, 1c, and 2 constructs. RLU: relative luminescence unit. (*n* = 4 biologically independent samples). **f**, Immunoblot of indicated proteins in HEK-293T cells transfected with siRNAs to UBXD8 or p97 and indicated 2X-FLAG-tagged mature SREBP1a, 1c, and 2 constructs. Immunoblots were probed with antibodies to SREBP1 and 2 to visualize immature and transfected mature forms. **g**, Split luciferase assay to measure contacts in HEK 293T cells treated with SCD1 inhibitor MF438 at 1 *μ*M for 4 hours. Cells were also treated with oleic acid for 4 hours as indicated. (*n* ≥ 3 biologically independent samples). **h,** Immunoblot of indicated proteins in HEK293T cells transfected with indicated siRNAs and wildtype or catalytically dead mutant of SCD1. GFP- HA/FLAG was transfected as a negative control. Related for Fig. 3e. (*n* = 3 independent biological replicates). Data are means ± SEM (*, **, *** *P* < 0.05, 0.01, 0.0001 respectively. One-way ANOVA with Dunnett’s multiple comparison test (**b, c**), Paired *t* test with Welch’s correction (**d**) or One- way ANOVA with Tukey’s multiple comparison test (**e & g**).

**Supplementary Fig. 6.**
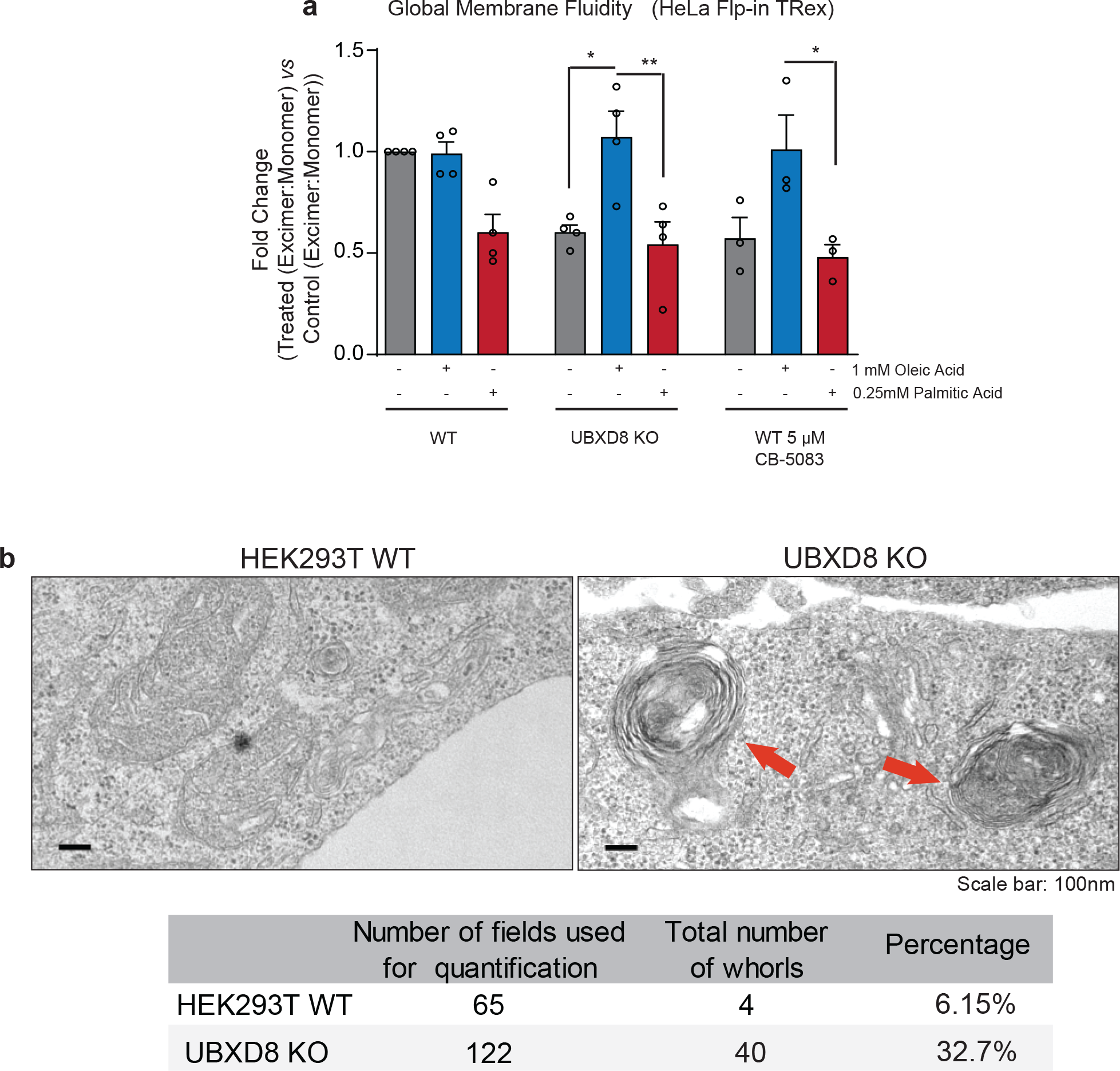
Loss of UBXD8 perturbs membrane saturation. **a,** Global membrane fluidity in cells was measured using pyrene-based lipid probes in wildtype and UBXD8 KO HeLa-Flp-IN-TRex cells. Wildtype cells were also treated with 5 *μ*M of the p97 inhibitor CB-5083 for 4 hours. Cells were supplemented with indicated concentrations of oleic acid and palmitic acid for 4 hours. The fold change (Treatedexcimer:monomer *vs* Controlexcimer:monomer) of ratio of excimer (Em. Max. 460nm) to monomer (Em max. 400nm) fluorescence is indicated. Fold changes < 1 indicate more ordered lipid bilayers relative to WT untreated control. (*n* = 3 biologically independent samples). **b**, Representative transmission EM micrographs of multilamellar bodies (red arrows) containing membrane whorls in UBXD8 KO HEK-293T cells. Lower panel shows quantification of multilamellar bodies from images in (**c**). (Measurements are from *n* = 3 biological replicates with WT = 50 cells in 65 fields and UBXD8 KO = 120 cells in 122 fields). Data are means ± SEM (*, **, \*\*\**P* < 0.05, 0.01, 0.0001 respectively, One-way ANOVA with Tukey’s multiple comparison test (**a**). Scale bar, 100 nm.

## Notes

### Competing Interest Statement

The authors have declared no competing interest.

